# Control of osteocyte dendrite formation by Sp7 and its target gene osteocrin

**DOI:** 10.1101/2021.03.22.436056

**Authors:** Jialiang S. Wang, Tushar Kamath, Fatemeh Mirzamohammadi, Daniel Rotter, Hironori Hojo, Christian D. Castro, Rushi Patel, Nicolas Govea, Tetsuya Enishi, Yunshu Wu, Janaina da Silva Martins, Michael Bruce, Daniel J. Brooks, Mary L. Bouxsein, Danielle Tokarz, Charles P. Lin, Abdul Abdul, Evan Z. Macosko, Melissa Fiscaletti, Craig F. Munns, Makoto Fujiwara, Henry M. Kronenberg, Marc N. Wein

**Author notes:** Address correspondence to: Marc N. Wein, Thier Research Building Room 1101, Endocrine Unit, Massachusetts General Hospital, Boston, MA 02114 USA.

## Abstract

Osteocytes use an elaborate network of dendritic connections to control bone remodeling. Some osteoblasts embed within mineralized bone matrix, change shape, and become osteocytes. The molecular circuitry that drives dendrite formation during “osteocytogenesis” is poorly understood. Here we show that deletion of *Sp7*, a gene linked to rare and common skeletal disease, in mature osteoblasts and osteocytes causes severe defects in osteocyte dendrites. Unbiased profiling of Sp7 target genes and binding sites reveals unexpected repurposing of this transcription factor to drive dendrite formation. *Osteocrin* is a Sp7 target gene that promotes osteocyte dendrite formation and rescues phenotypic and molecular defects in Sp7-deficient mice. Single-cell RNA-sequencing demonstrates overt defects in osteocyte maturation *in vivo* in the absence of Sp7. Sp7-dependent gene networks enriched in developing osteocytes are associated with rare and common human skeletal traits. Moreover, humans homozygous for the osteogenesis imperfecta-causing *SP7^R316C^* mutation show dramatic defects in osteocyte morphology. Genes that mark osteocytes *in vivo* and that are regulated by Sp7 *in vitro* are highly enriched in neurons, highlighting shared features between osteocytic and neuronal connectivity. Taken together, these findings reveal a crucial role for Sp7 and its target gene *Osteocrin* in osteocytogenesis, demonstrating that pathways that control osteocyte development influence human bone diseases.

## Introduction

The major cell types that govern bone homeostasis are osteoblasts, osteoclasts, and osteocytes. While the roles of osteoblasts and osteoclasts in bone formation and resorption have been well studied [1, 2], those of osteocytes, the most abundant cell type in bone, had been overlooked due to technological limitations and the cells’ relatively inaccessible location within mineralized bone matrix. Recently, emerging evidence has highlighted key roles for osteocytes in bone remodeling [3]. These cells translate external cues, such as hormonal variations and mechanical stresses, into changes in bone remodeling by secreting paracrine-acting factors that regulate osteoblast and osteoclast activity [4]. Furthermore, osteocytes have a unique morphology as they bear multiple long, neuron-like dendritic processes projecting through the lacunar canalicular system in bone [5]. The osteocyte dendritic network confers mechano-sensitivity to these cells, and allows for extensive communication amongst osteocytes and adjacent cells on bone surfaces [6]. Defects in the osteocyte dendrite network may cause skeletal fragility in the setting of aging and glucocorticoid treatment [7, 8]. Recent estimates suggest that the osteocyte connectivity network in human bone exhibits the same order of complexity as the network of connections between neurons in the brain [9].

Lineage-specifying transcription factors have been identified for other key cell types in bone [10]; however, lineage-defining transcription factors that coordinate genetic programs associated with osteocyte maturation remain unknown. The goal of this study was to define key gene regulatory circuits that drive osteocyte differentiation and dendrite formation. Osterix/Sp7 (encoded by the *Sp7* gene) is a zinc finger-containing transcription factor essential for osteoblast differentiation and bone formation downstream of Runx2 [11]. Common human *SP7*-associated variants are linked to bone mineral density variation and fracture risk [12, 13], and rare *SP7* mutations cause recessive forms of osteogenesis imperfecta [14, 15]. Here, we deleted Sp7 in mature osteoblasts and osteocytes and observed a dramatic skeletal phenotype including cortical porosity, increased osteocyte apoptosis, and severe defects in osteocyte dendrites. Unbiased profiling of Sp7 target genes and binding sites in osteocytes revealed a novel stage-specific role for this transcription factor during osteocytogenesis. Osteocyte-specific Sp7 target genes have DNA binding sites distinct from those in osteoblasts and are highly enriched in genes expressed in neurons, highlighting novel shared molecular links between inter-cellular communication networks in brain and bone. Amongst osteocyte-specific Sp7 target genes, we identify osteocrin as a secreted factor that promotes osteocyte dendrite formation/maintenance *in vitro* and *in vivo.* Single-cell RNA-sequencing identified discrete populations of cells undergoing the osteoblast-to-osteocyte transition, and dramatic defects in this normal process in the absence of Sp7. Finally, we report overt defects in osteocyte dendrites in humans with bone fragility due to the rare *SP7^R316C^* mutation. Taken together, these findings highlight a central role for Sp7 in driving cell morphogenesis during osteocytogenesis.

## Results

### Sp7 deletion in mature osteoblasts leads to severe osteocyte dendrite defects

We sought to identify gene regulatory networks that control osteocytogenesis [16, 17]. *Sost* is uniquely expressed in osteocytes but not osteoblasts [18, 19], so we hypothesized that transcription factors that drive *Sost* expression will control other aspects of osteocyte function. Combined over-expression of *Atf3*, *Klf4*, *Pax4*, and *Sp7* induced ectopic *Sost* expression [20]. Therefore, we asked if these transcription factors might participate in eutopic *Sost* expression in Ocy454 cells [21]. Of these factors, only *Sp7* controlled sclerostin secretion (Supplementary Fig. 1a). To examine the role of Sp7 in osteocytes *in vivo*, we deleted *Sp7* using *Dmp1-Cre* [22], which targets mature osteoblasts and osteocytes (Fig. 1a, e). *Sp7^OcyKO^* mice showed increased cortical porosity (Fig. 1b, e), reduced cortical bone mineral density (Fig. 1e and Supplementary Table 1a), and abnormal intracortical bone remodeling as characterized by increased intracortical bone formation and resorption (Fig. 1c-d, Supplementary Table 1b).

**Fig. 1:**
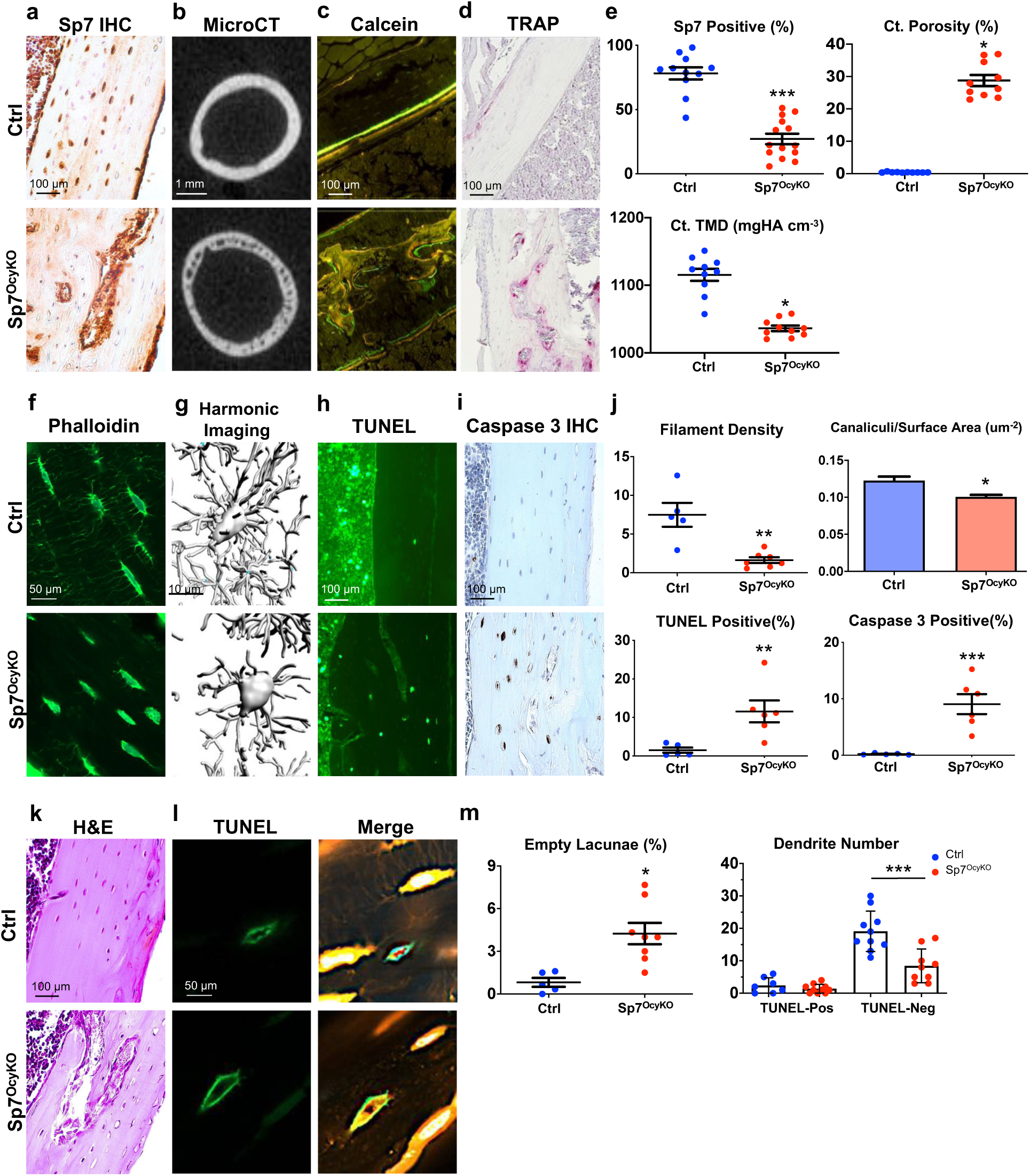
Severe skeletal defects in *Sp7^OcyKO^* mice. (**a**) Sp7 immunohistochemistry was performed on tibiae from 8-week-old control (*Dmp1-Cre; Sp7^+/+^*) and *Sp7^OcyKO^* (*Dmp1-Cre; Sp7^f/f^*) mice, quantification of Sp7-positive osteocytes is shown in (**e**). (**b**) Cross sectional µ-CT images from the femoral midshaft diaphysis reveal reduced mineralization and increased cortical porosity in *Sp7^OcyKO^* mice, quantified in (**e**). (**c**) 8-week-old mice were labeled with calcein (green) and demeclocycline (red) 7 and 2 days prior to sacrifice, respectively. Non-decalcified sections from cortical bone in the tibia were analyzed. Control mice show orderly endosteal bone formation. In contrast, *Sp7^OcyKO^* animals show abnormal intracortical bone formation. (**d**) TRAP-stained (red) paraffin-embedded sections from the tibia show increased number of intra-cortical osteoclasts *Sp7^OcyKO^* in mice. (**f**) Cryosections from 8-week-old control (*Dmp1-Cre; Sp7^+/+^*) and *Sp7^OcyKO^* (*Dmp1-Cre; Sp7^f/f^*) tibia were stained with phalloidin to visualize actin filaments, panel (**j**) shows quantification indicating reduced filament density (percent of acellular bone matrix occupied by phalloidin-positive filaments) in *Sp7^OcyKO^* mice. (**g**) *In vivo* third generation harmonic imaging of the skull was performed to visualize osteocyte cell bodies and canaliculi in the skull. Quantification of defects in canaliculi/surface area is shown in (**j**). (**h-j**) Apoptosis *in situ* was analyzed on tibia sections by TUNEL and activated caspase 3 immunohistochemistry; both methods demonstrate increased osteocyte apoptosis in *Sp7^OcyKO^* mice versus controls. (**k**) Hematoxylin and eosin-stained paraffin-embedded sections from the tibia show abnormal cortical porosity and empty osteocyte lacunae, quantified in (**m**). (**l-m**) TUNEL staining (green) was performed on control (*Dmp1-Cre; Sp7^+/+^, Ai14*) and *Sp7^OcyKO^* (*Dmp1-Cre; Sp7^f/f^, Ai14*) mice. Thereafter, osteocyte filaments were visualized by tdTomato fluorescence within dendritic projections. Dendrite number was counted in TUNEL positive (rare in control mice) and TUNEL negative osteocytes. Reduced dendrite numbers are noted in non-apoptotic (TUNEL-negative) osteocytes lacking Sp7. *, p<0.05, **, p<0.01. ***, p<0.001, ****, p<0.0001.

*In situ* phalloidin staining of actin filaments and *in vivo* third harmonic generation (THG) microscopy [23] both revealed dramatic reductions in osteocyte dendrites in *Sp7* mutants compared to controls (Fig. 1f, g and j; Supplementary Fig. 1b). Osteocyte morphology defects seen here are similar to those previously reported in global, inducible *Sp7* mutant mice [24]. Osteocyte apoptosis was elevated in *Sp7^OcyKO^* mice (Fig. 1h, i and j), and *Sp7* conditional mutants showed increased numbers of empty lacunae (Fig. 1k, m). Dendrite numbers were reduced even in non-apoptotic *Sp7^OcyKO^* osteocytes (Fig. 1l-m), suggesting that apoptosis may not be the cause of decreased dendrite number. These results indicate that Sp7, a transcription factor crucial for early steps in osteoblast lineage commitment [11], continues to play a key role in maintaining skeletal integrity throughout the osteoblast/osteocyte lineage where it is required for normal dendritic morphology. Without Sp7, dysmorphic osteocytes undergo apoptosis which triggers subsequent intracortical remodeling [25] and gross defects in cortical bone integrity.

### Sp7 deficiency *in vitro* impedes dendrite formation

While *Sp7* knockdown caused no obvious morphology differences in normal tissue culture conditions (Fig. 2a), cells lacking Sp7 showed reduced dendrite numbers compared to control cells (Fig. 2a-b; Supplementary Fig. 1d) grown in 3D collagen gels. Transcriptomic analysis of 2D versus 3D cultured cells revealed coordinate changes in 3D culture consistent with osteocyte maturation (Supplemental Fig. 2c and Supplemental Table 2). *Sp7* knockdown caused reduced cell numbers over time in culture (Fig. 2c), a defect that is accompanied by unchanged proliferation (Fig. 2d) and, consistent with our *in vivo* findings (Fig. 1h-i), increased apoptosis (Fig. 2d). These findings support the utility of this 3D culture system for studies on the role of Sp7 in osteocytogenesis, and demonstrate that Sp7 is necessary for osteoblasts to undergo an osteocyte-like morphology change.

**Fig. 2:**
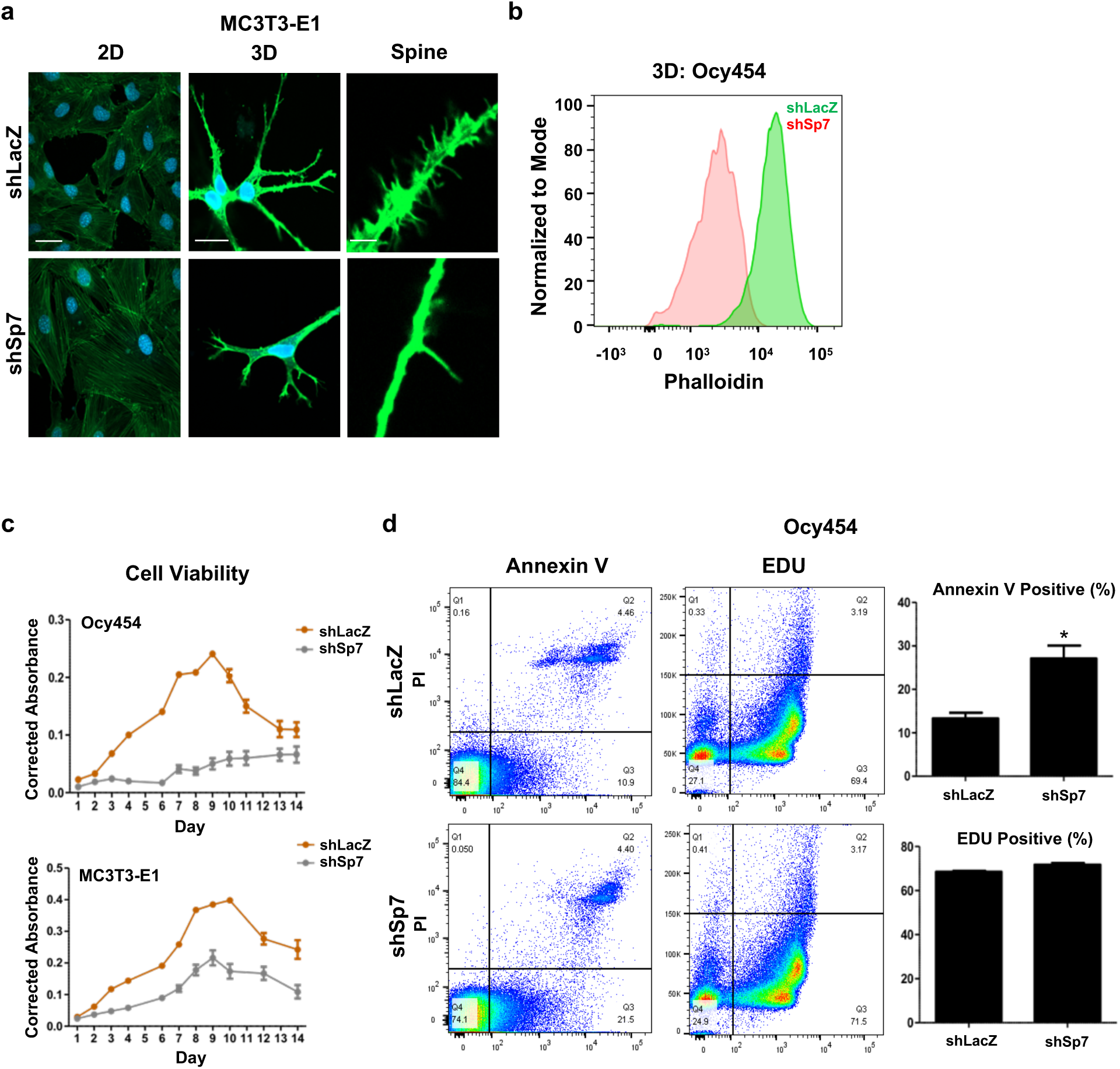
Sp7 is required for optimal dendrite formation *in vitro*. (**a**) Control (shLacZ) and *Sp7* knockdown (shSp7) MC3T3-E1 cells were cultured in standard (2D) or 3D (type I collagen gel) conditions, then stained with phalloidin (green) and DAPI (blue). In 3D culture, *Sp7* knockdown cells show short dendrites (middle panel) with reduced complexity (right panel). Scale bars for 2D and 3D images represent 10 µm, scale bar for “spine” images represent 1 µm. (**b**) Control and *Sp7* knockdown Ocy454 cells were grown in 3D culture then stained with FITC-Phalloidin for flow cytometry. Reduced intracellular phalloidin staining is noted in *Sp7* knockdown cells. (**c**) Growth curves for control and *Sp7* knockdown Ocy454 cells (top) and MC3T3-E1 cells (bottom) were determined using a resavurin-based viability dye. (**d**) Control and *Sp7* knockdown Ocy454 cells were analyzed *in vitro* for apoptosis and proliferation. Normal proliferation is noted based on EdU incorporation, while *Sp7* knockdown cells show increased apoptosis.

### Osteocyte Sp7 target genes are involved in dendrite formation

The osteocyte network in bone bears similarity to the network of inter-cellular connections between neurons [9]. In addition, osteocytes have been reported to express certain neuronal-enriched transcripts [26]; however, the molecular mechanisms used by osteocytes to acquire this gene expression program are unknown. RNA-seq in Sp7 deficient and over-expressing Ocy454 cells (Fig. 3a-d; Supplementary Fig. 2a; Supplementary Table 2) revealed that Sp7-dependent genes are enriched in gene ontology terms linked to cell projection organization and neuronal development.

**Fig. 3:**
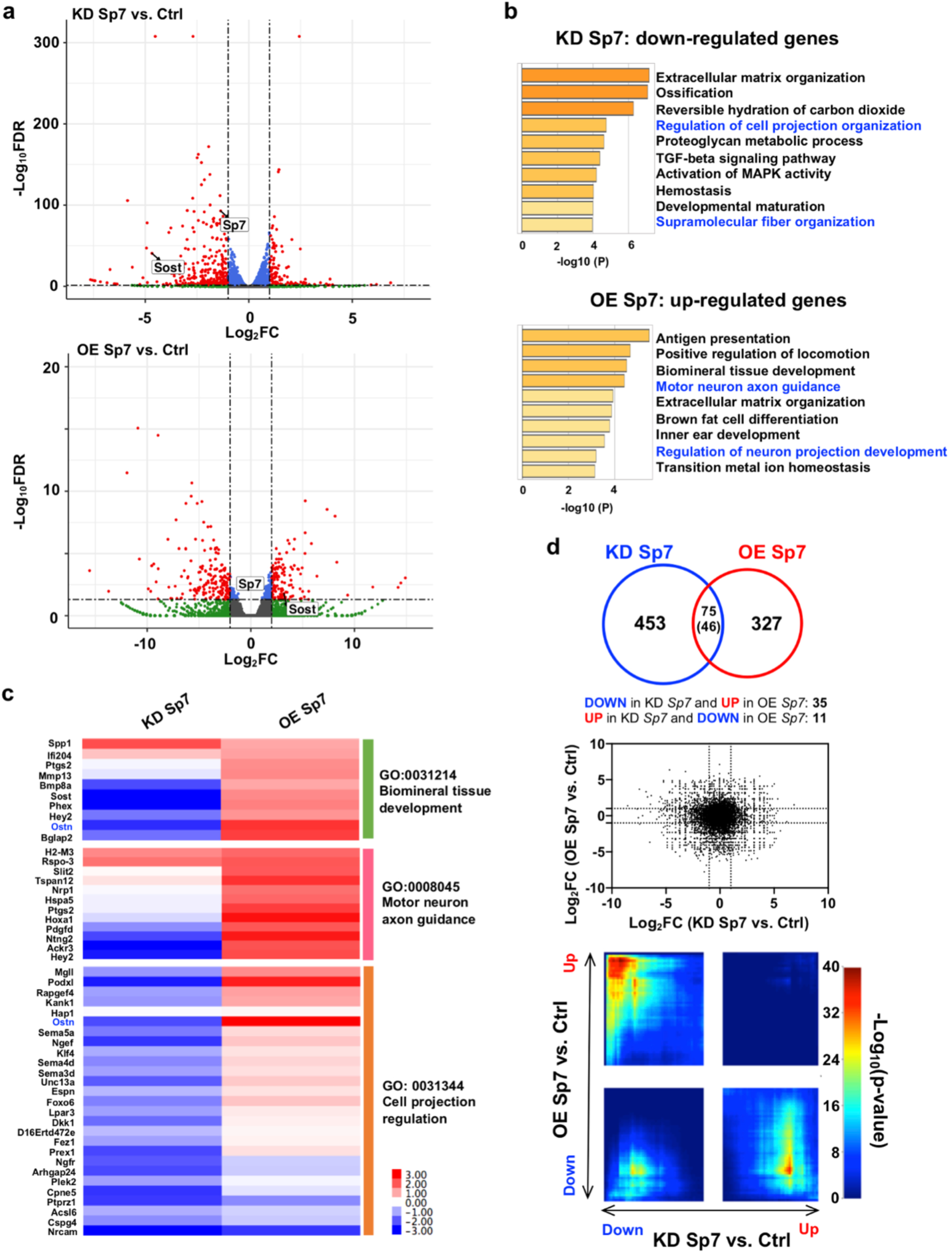
Sp7 controls expression of genes involved in neuron projection development in osteocytes. (**a**) RNA-seq was performed on control (shLacZ) and *Sp7* knockdown (KD, top) and control (LV-GFP) and *Sp7* overexpressing (OE, bottom) Ocy454 cells. Volcano plots demonstrate differentially-expressed genes (red data points) in each comparison. See Supplementary Table 2. (**b**) Gene ontology analysis of genes down-regulated by *Sp7* knockdown (top) and up-regulated by *Sp7* overexpression (bottom) reveals enrichment in several terms associated with cell projection development and neuronal morphogenesis. (**c**) Heatmap showing fold change regulation of individual genes in gene ontology groups of interest. (**d**) Relationship between gene expression changes in response to Sp7 perturbation. Top, Venn diagram revealing number of genes regulated by both *Sp7* overexpression and *Sp7* knockdown. Middle, scatterplot showing fold change regulation of all detected genes by *Sp7* knockdown (x-axis) and *Sp7* overexpression (y-axis). Bottom, RRHO2 visualization revealing statistically significant groups of genes counter-regulated by *Sp7* knockdown versus overexpression.

Next, we performed Sp7 ChIP-seq in Ocy454 cells (Fig. 4a; Supplementary Table 3; studies were performed in conditionally-immortalized Ocy454 cells due to challenges obtaining sufficient numbers of primary osteocytes for ChIP-seq) and compared this dataset to the published Sp7 cistrome in primary osteoblasts (POB) [27] to define osteocyte-specific (Ocy-specific), primary osteoblast-specific (POB-specific), and shared Sp7 (Ocy-POB) enhancer binding sites (Fig. 4a-b). Ocy-specific Sp7 enhancer peaks are enriched in genes associated with axon guidance (Fig. 4c, left) and Ocy-specific Sp7 promoters are enriched in genes associated with actin filament assembly (Fig. 4c, right). Systematic comparison of the chromatin landscape [28] at Sp7-bound Ocy-specific enhancer sites revealed that histone modifications surrounding Sp7-bound Ocy-specific enhancers were in largely the same state in both osteocytes and primary osteoblasts (Fig. 4d). Therefore, Sp7 does not globally change the epigenetic state during the osteoblast-to-osteocyte transition, but rather acts upon open regulatory regions shared between those two cell types (Fig. 4e).

**Fig. 4:**
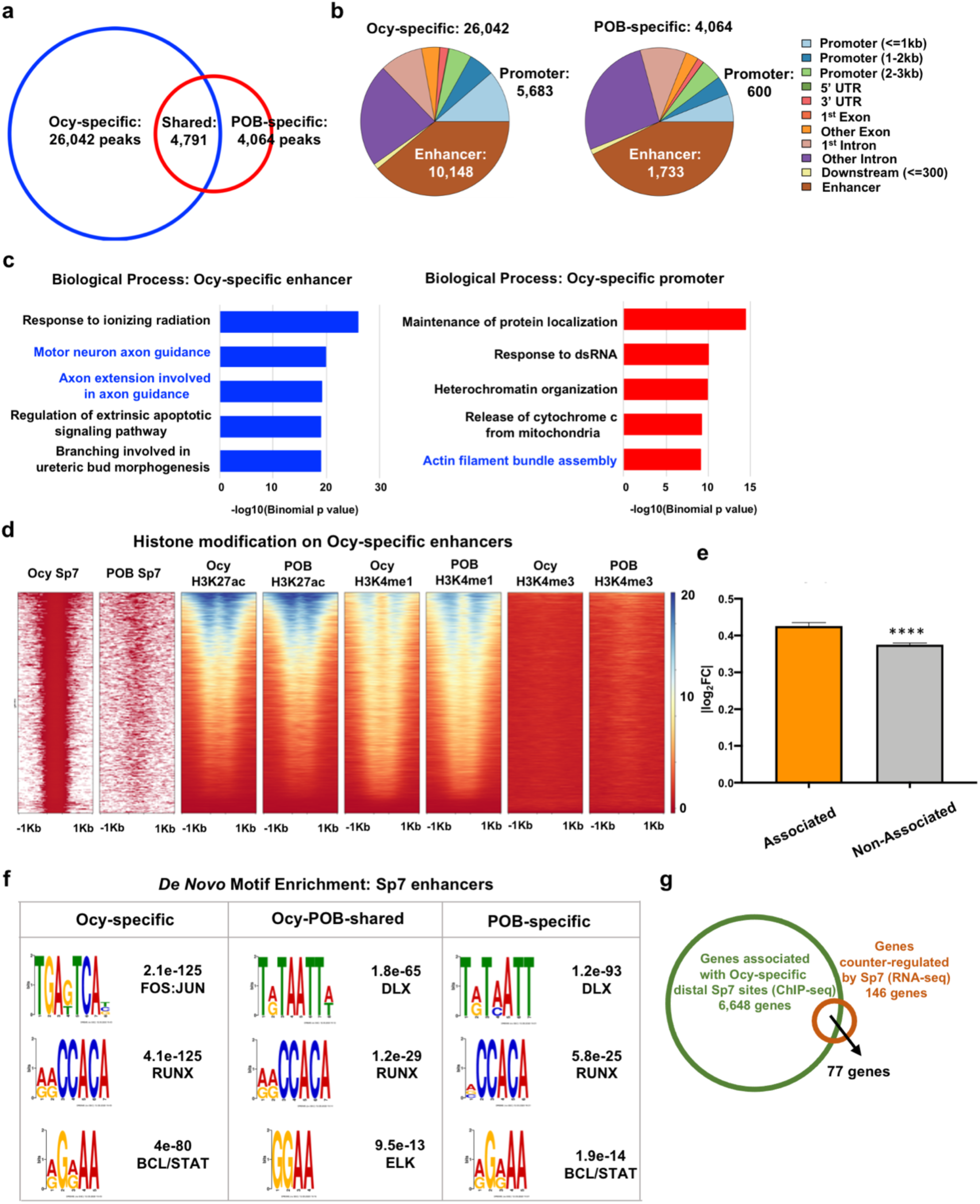
Definition of the osteocyte-specific Sp7 cistrome. (**a**) Sp7 ChIP-seq was performed in Ocy454 cells, and Sp7 binding patterns were compared between Ocy454 cells and primary osteoblasts (POB). See also Supplementary Table 3. (**b**) Genomic distribution of cell-type specific Sp7 binding sites. (**c**) Gene ontology analysis of genes linked to Ocy454 cell-specific Sp7 peaks. Sp7 associates with regulatory regions of genes linked to motor neuron axon guidance and actin filament bundle assembly. (**d**) Ocy454 cell-specific Sp7 bound enhancer peaks were analyzed for the indicated histone modifications in d3 (POB) and d35 (Ocy) IDG-SW3 cells. See text for details. (**e**) Genes were categorized based on the presence of Ocy-specific Sp7 enhancer association in Ocy454 cells. Then, the expression of genes in Sp7 shRNA versus LacZ shRNA was reported as the absolute value of the log2 fold change. Genes bound by Sp7 show, on average, greater effect of *Sp7* knockdown than genes without Sp7 enhancer binding sites. (**f**) *De novo* motif analysis of osteocyte-specific, osteoblast-specific, and shared distal Sp7 peaks. See also Supplementary Table 4. (**g**) 77 genes (Supplementary Table 5) are revealed by intersecting transcripts regulated by both *Sp7* knockdown and overexpression in Ocy454 cells (FDR < 0.05, 146 genes) and genes with associated Ocy454 cell specific Sp7 ChIP-seq peaks (6,648 genes). ****, p<0.0001

In osteoblasts, Sp7 binds enhancer chromatin indirectly via associating with Dlx family transcription factors [27]; therefore, we asked if Sp7 might utilize a distinct binding cofactor in osteocytes. Comparison of enriched sequence motifs present in each group of enhancers [29] demonstrated that Ocy-specific, shared, and POB-specific enhancers all showed enrichment of binding motifs associated with skeletal development (Supplementary Table 4). As expected [27], POB-specific enhancers are enriched in Dlx binding sequences (Fig. 4f). In contrast, Ocy-specific enhancers demonstrated selective enrichment for the TGA(G/T)TCA motif bound by AP1 family members (Fig. 4f) [30, 31]. Therefore, Sp7 likely cooperates with distinct DNA binding transcription factors to regulate enhancer activity in osteoblasts versus osteocytes.

We next intersected RNA-seq and ChIP-seq datasets to identify 77 direct Sp7 target genes in osteocytes (Fig. 4g; Supplementary Figure 2c; Supplementary Table 5). The relative mean expression values of these 77 genes, and a distinct group of 134 POB-specific Sp7 target genes, are significantly enriched in neurons versus other cell types in mouse brain [32] (Supplementary Figure 7b-c). This observation demonstrates strong convergence between bone cell Sp7 target genes and neuronally-expressed genes.

### Osteocrin rescues morphologic and molecular defects caused by Sp7 deficiency

Osteocyte-specific Sp7 targets were selected for subsequent functional studies. CRISPR/Cas9-mediated deletion of the secreted peptide Osteocrin (encoded by the *Ostn* gene) [33] selectively reduced phalloidin staining in 3D culture (Supplementary Fig. 3a-b). The *Ostn* gene encodes a small secreted protein that is expressed is periosteal cells in mouse and human bone [33, 34] (Supplementary Fig. 3c shows higher level of periosteal *Ostn* expression in control compared to Sp7 mutant bones), muscle, and primate neurons that potentiates C-type Natriuretic Peptide (CNP) signaling [35, 36] (Supplementary Fig. 3d).

Since *Ostn* is an osteocyte-specific Sp7 target gene whose levels are reduced in *Sp7* knockdown cells (Fig. 3c), we asked if *Ostn* over-expression could rescue defects associated with Sp7 deficiency *in vitro*. *Ostn* over-expression in *Sp7* knockdown cells led to rescue of dendrite numbers and F-actin content in 3D culture (Fig. 5a). Next, we used RNA-seq (Supplementary Fig. 4a-b; Supplementary Table 6) where we predicted that genes whose expression was dependent on Sp7 might be rescued by *Ostn* over-expression. Indeed, 516 out of 560 (92.1%) of the genes differentially regulated in both comparisons (shLacZ vs shSp7 and shSp7: control vs osteocrin overexpression) were counter-regulated by Sp7 and Ostn (Supplementary Fig. 4c-d). Scatterplot analysis and RRHO2 threshold test demonstrated strong inverse correlation between profiles seen with *Sp7* knockdown and *Ostn* rescue (Supplementary Fig. 4c; Fig. 5b).

**Fig. 5:**
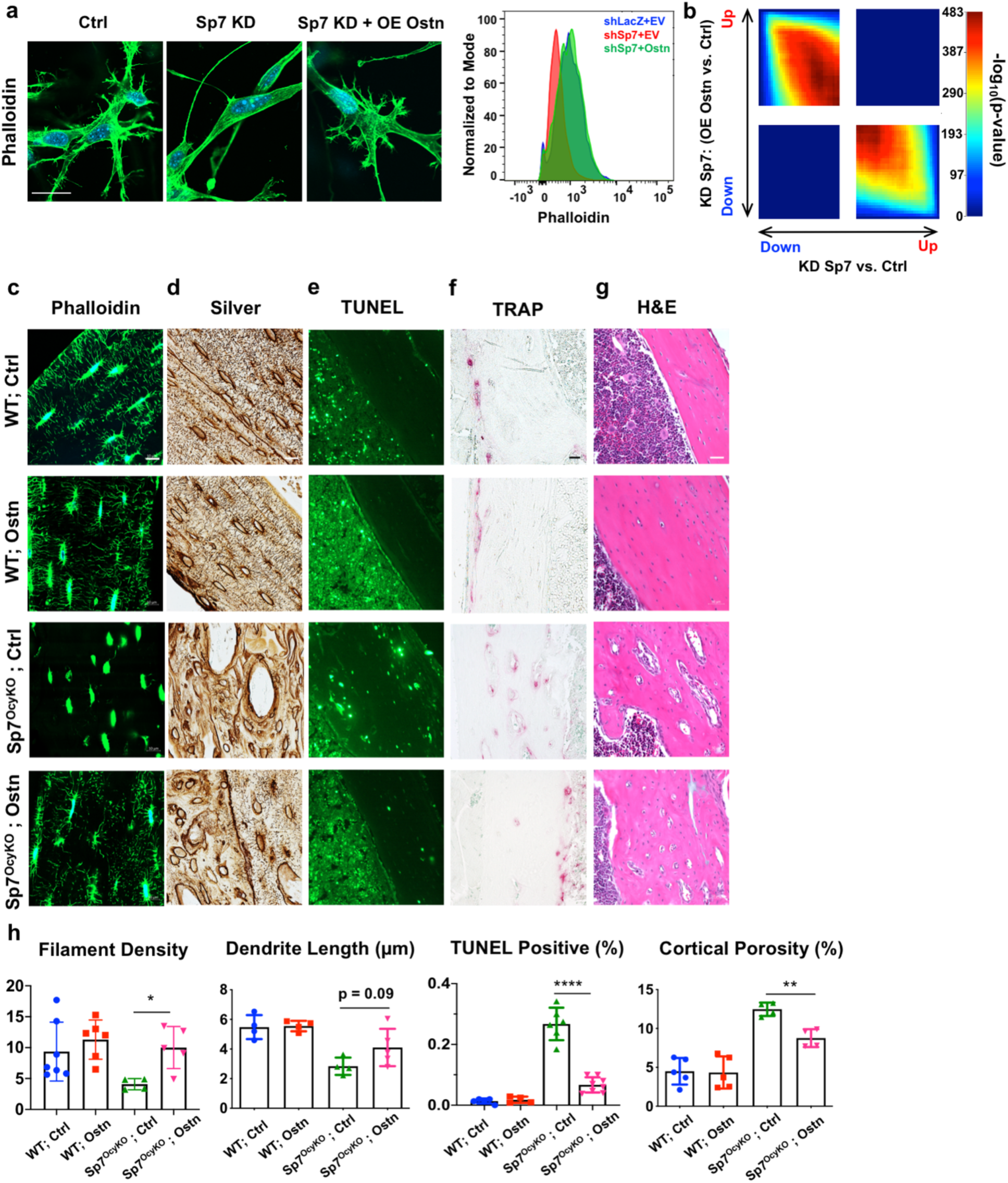
Exogenous osteocrin rescues skeletal phenotypes associated with Sp7 deficiency. (**a**) *Sp7* knockdown (KD) MC3T3-E1 cells were infected with control (empty vector) or osteocrin (LV-Ostn) lentiviruses followed by growth in 3D collagen gels and phalloidin (green) and DAPI (blue) staining. *Osteocrin* overexpression rescues phalloidin staining intensity to levels observed in control (shLacZ) cells. Scale bar represents 10 µm. (**b**) RRHO2 visualization revealing global counter-regulation of gene sets by *Sp7* knockdown and osteocrin rescue. (**c-h**) Control or *Sp7^OcyKO^* mice were injected with AAV8-control or AAV8-osteocrin viral particles at 3 weeks of age, then analyzed 3 weeks later for histologic analysis by phalloidin staining (**c**), silver staining (**d**), TUNEL (**e**), TRAP (**f**), or H & E (**g**). The scale bar in (**c-d**) is 10 µm. The scale bar in (**e-g**) is 50 µm. All results are quantified in panel (**h**).

Next, we asked if increasing systemic Ostn levels might rescue osteocyte defects observed in *Sp7* mutant mice. To do this, we treated *Sp7^OcyKO^* mice with an *Ostn*-expressing adeno-associated virus (AAV8, a strategy that increases circulating Ostn levels by boosting hepatic Ostn secretion) (Supplementary Fig. 4e). AAV8-Ostn rescued reduced dendrite number, dendrite length, and osteocyte apoptosis in *Sp7^OcyKO^* mice (Fig. 5c-e, h). These AAV8-Ostn effects on osteocyte morphology and survival were associated with reduced intracortical osteoclasts and reduced cortical porosity (Fig. 5f-h). In sum, over-expression of the Sp7 target gene *Ostn* rescues the morphologic, transcriptomic, and phenotypic defects seen when Sp7 is absent in mature osteoblasts and osteocytes.

### Sp7-dependent transitional cells during osteocyte development revealed by single-cell transcriptomics

The discrete transitional cell types between bone-forming osteoblasts and matrix-embedded, dendrite-bearing osteocytes are currently unknown. We used *Dmp1*-*Cre*; tdTomato reporter mice and adapted an aggressive digestion protocol to liberate these cells from bone matrix. Viable tdTomato^+^ bone cells from control and *Sp7^OcyKO^* were recovered (see Methods) for high-throughput single-cell RNA-seq (scRNA-seq) (Fig. 6, Supplementary Fig. 5 and 6). Using LIGER [37], we captured eight distinct *Bglap3*-expressing “Ob/Ocy” osteo-lineage clusters, ranging from proliferating pre-osteoblasts to mature *Sost*-expressing osteocytes (Fig. 6a; Supplementary Fig. 6; Supplemental Table 7) [37]. Based upon expression of enriched genes in each cluster, we annotated these 8 groups of cells as (1) *Pdzd2/Bglap3* (canonical osteoblasts), (2) *Pdgfrl/Mgst3*, (3) *Tnc/Enpp1*, (4) *Plxdc2/Kcnk2*, (5) *Tagln2/Dpysl3*, (6) *Fbln7/Sost* (mature osteocytes), (7) *Adipoq/Cxcl12* (CAR cells), and (8) *Mki67* (proliferating pre-osteoblasts). *Tnc*, *Kcnk2* and *Dpysl3* mark 3 intermediate subpopulations (clusters 3, 4, and 5, respectively, between osteoblasts and osteocytes (Fig. 6b). *Tnc* and *Dpysl3* are highly expressed in subpopulations of cells in the endosteum and periosteum, and in some early mineralizing osteocytes near bone surfaces (Fig. 6c). *Fbln7* is expressed in newly-formed osteocytes and mature osteocytes buried deep within bone matrix (Fig. 6c).

**Fig. 6:**
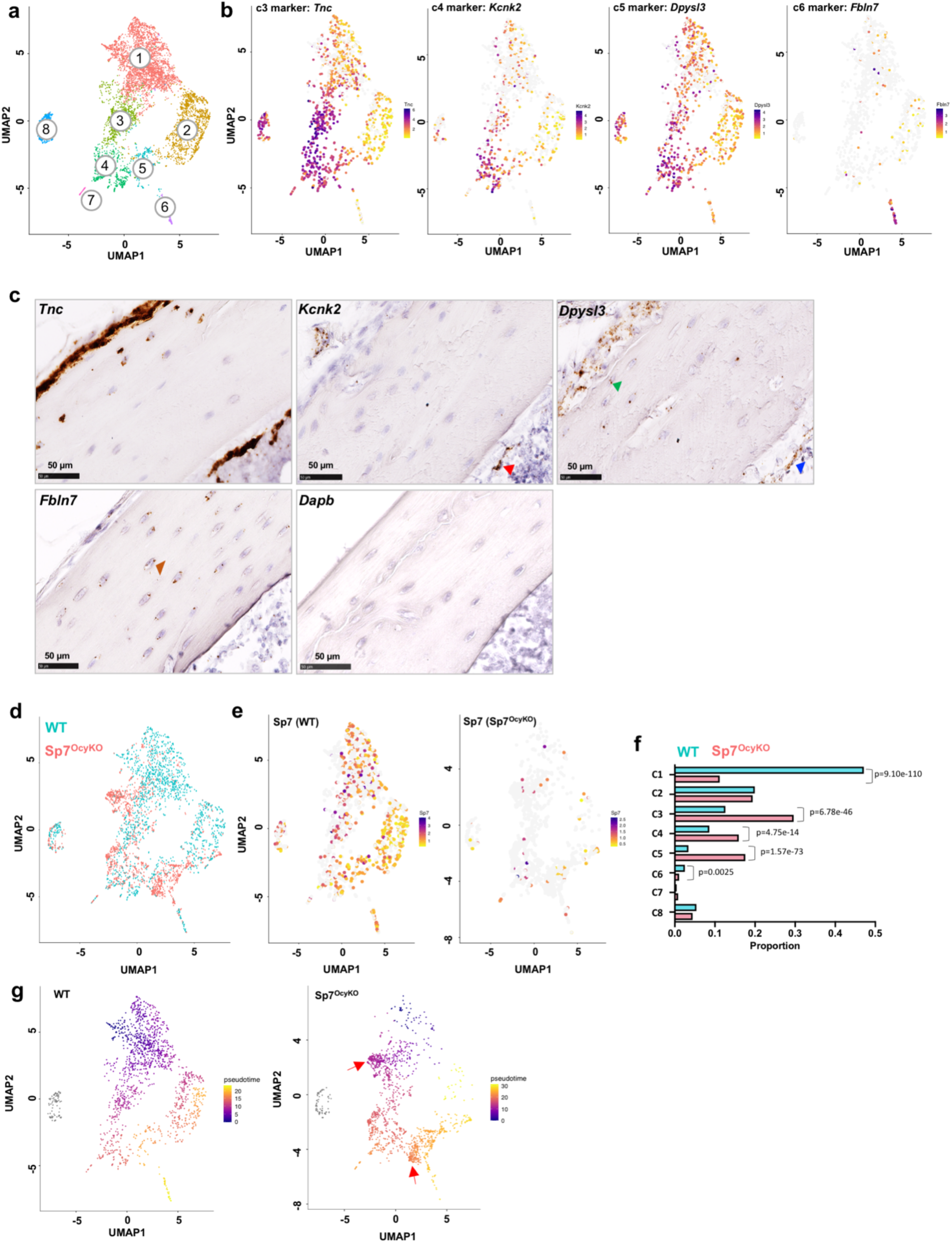
Single cell transcriptomic profiling of mature osteoblasts and differentiating osteocytes highlights a key role for Sp7. (**a**) Long bones were subjected to serial collagenase/EDTA digestions (see methods), and cells from fractions 5, 7, and 8 were collected for flow cytometry. Viable (DAPI-negative) tdTomato-positive cells were sorted followed by single-cell RNA-seq library construction. See also Supplementary Fig. 5 and 6. Cell-clustering from WT mice was performed. (**b**) Feature plots showing the expression of cluster-specific markers: *Tnc* (c3); *Kcnk2* (c4); *Dpysl3* (c5); *Fbln7* (c6). (**c**) RNA *in situ* hybridization of cluster-specific markers: *Tnc, Kcnk2, Dpysl3* and *Fbln7*. *Kcnk2* expression (red triangle) is primarily noted in endosteal cells that form a canopy around osteoblast. *Dpysl3* expression is noted in osteocytes close to bone surfaces (green) and a subpopulation of endosteal cells (blue). *Fbln7* expression is noted only in embedded osteocytes (brown). Negative control: *Dabp*. (**d**) Integrated analysis of cells from WT (blue) and *Sp7^OcyKO^* (red) mice after down-sampling of WT library to match cellular representation seen in *Sp7* mutant mice. (**e**) Feature plot showing *Sp7* expression across 8 Ob/Ocy clusters. As expected, *Sp7* expression is reduced in mutant mice. (**f**) Relative proportions (out of 1.0) in WT vs *Sp7^OcyKO^* mice. *Sp7* mutants show reduced canonical osteoblasts (cluster 1), increased cells in clusters 3-5, and reduced cells in cluster 6 (mature osteocytes). (**g**) Monocle3 analysis showing pseudo-time trajectory mapping across tdTomato-positive cells in WT (left) and *Sp7^OcyKO^* (right) mice. Red arrows indicate Sp7-specific subpopulations in clusters 3 and 5 demonstrating apparent arrested differentiation.

Comparison of tdTomato^+^ cells between *Sp7^OcyKO^* mice and littermate controls revealed striking changes in cluster cellularity in *Sp7^OcyKO^* mice (Fig. 6d-f). *Sp7* mutant mice showed increased relative proportions of cells in transitional clusters 3, 4, and 5 and reduced numbers of mature osteocytes, further demonstrating arrested osteocyte maturation in the absence of Sp7. Next, we performed pseudo-time differentiation analysis using Monocle3 [38] and Velocyto [39] (Fig. 6g; Supplementary Fig. 5f). These analyses suggest the potential of two differentiation pathways from canonical osteoblasts (cluster 1) via intermediate cells in clusters 3 > 4 > 5 or 2 > 5 into mature *Fbln7/Sost*-expressing osteocytes (cluster 6). Compared to the orderly trajectory seen in WT cells, *Sp7* mutants show multiple defects (red arrows in Fig. 6g) including apparent arrest in clusters 3 (marked by *Plec*) and 5 (marked by *Lmo2*). Taken together, these single-cell RNA-sequencing results confirm the key role for Sp7 in orchestrating multiple steps in osteocyte differentiation *in vivo*.

Cluster-specific differential gene expression analysis in intermediate clusters (Supplementary Fig. 5e) revealed dysregulated mineralization-related genes in *Sp7^OcyKO^* cells in clusters 3, 4, and 5. Among dysregulated genes in *Sp7^OcyKO^* cluster 5 cells, *Lmo2* marks expression of a distinct subgroup of cells arrested in the transition to mature osteocytes (Supplementary Fig. 6 and see below). *Sp7^OcyKO^* cells show reduced *Pdpn* and *Fbln7* expression in cluster 6, consistent with blocked terminal osteocyte maturation (Supplementary Fig. 6). Furthermore, Sp7 deficiency is associated with marked dysregulation of terminal osteocyte markers (*Irx5*, *Ptprz1*) across the entire population of tdTomato+ cells captured. While these genes are normally restricted to mature osteocytes, they are aberrantly expressed in multiple sub-populations in *Sp7^OcyKO^* mice (Supplementary Fig. 6).

Gene Ontology analysis of the top 150 cluster 6 (*Fbln7/Sost*) osteocyte markers revealed enrichment with ‘neuronal’ terms such as cell projection organization and neuron differentiation (Supplementary Fig. 9a-b). To further explore potential similarity between osteocytes and neurons at the transcriptomic level, we performed enrichment analysis of top mature osteocyte markers across cell types in mouse brain (using methods detailed in Supplementary Fig. 7b, d). Osteocyte, but not osteoblast, marker genes are significantly enriched in their relative mean expression values in neurons versus other cell types in mouse brain (Supplementary Fig. 7d; Supplemental Fig. 8).

### Sp7 and osteocyte-linked genes are associated with common and rare human skeletal dysplasias

Next, we sought to further establish the relevance of the osteocyte functions of SP7 in human bone diseases. First, we used MAGMA [40] to understand the relationship between genes enriched in cells undergoing the osteoblast-to-osteocyte transition with genes linked to human BMD variation and fracture risk [13]. This analysis demonstrated significant enrichment of marker genes identified from osteoblasts and osteocytes, but not other tdTomato^+^ cells isolated and sequenced in our scRNA-seq analyses, with genes linked to human fracture risk (Fig. 7a). We then examined the enrichment of osteocyte markers in a recent classification of genes that when mutated cause distinct classes of human skeletal diseases [41]. The top osteocyte markers are significantly enriched in other groups of genes that cause ‘sclerosing bone disorders’ and ‘abnormal mineralization’ (Fig. 7b).

**Fig. 7:**
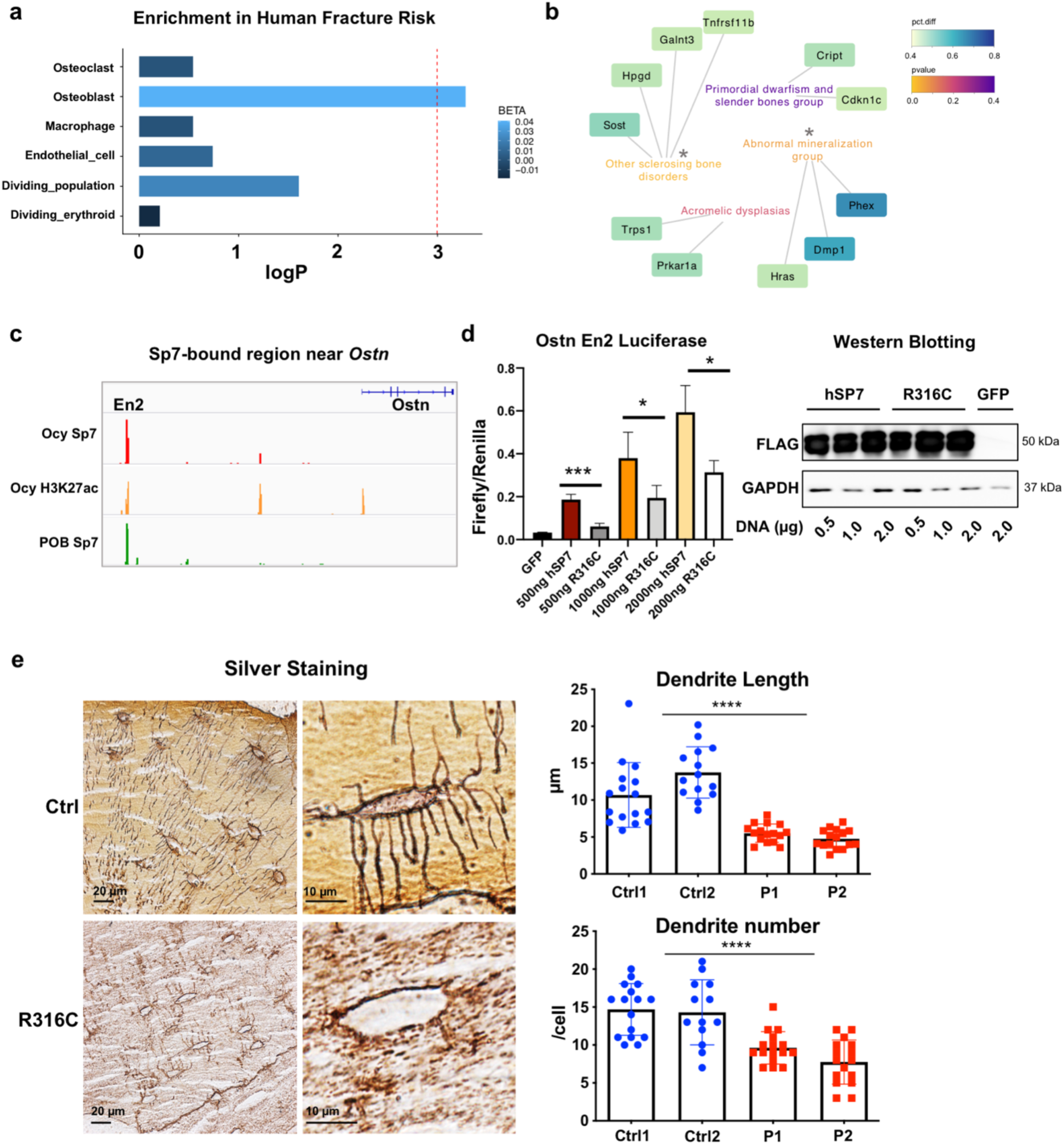
Sp7 and osteocyte-linked genes are associated with common and rare human skeletal diseases. (**a**) Negative log Bonferroni-corrected p-values obtained from the MAGMA gene-set analyses for all major cell types sampled from WT mice, shaded by effect size (BETA) values from the MAGMA gene-set regression. (**b**) Association between skeletal dysplasia disease groups and genes enriched in the top mature osteocyte markers. Two disease groups are significantly enriched: ‘Other sclerosing bone disorder’ (p = 0.016) and ‘Abnormal mineralization’ group (p = 0.039). (**c**) Locations of Sp7 and H3K27ac binding in the osteocrin regulatory region in Ocy454 cells and primary osteoblasts (POB). (**d**) Ostn_En2 activity is induced by WT, but not R316C, FLAG-tagged *SP7* over-expression in HEK293T cells. Comparable expression of both FLAG-tagged SP7 versions is noted by immunoblotting, with GAPDH as a loading control. (**e**) Non-decalcified iliac crest biopsy samples from age/sex-matched and *Sp7^R316C^* patient samples were silver stained to assess osteocyte morphology. Results are quantified in the right panel.

Patients homozygous for *SP7^R316C^* present with osteogenesis imperfecta-like bone disease including multiple fragility fractures and short stature [14, 15]. Sp7 binds an enhancer approximately 110 kB upstream of the *Ostn* transcription start site (Fig. 7c). The transcriptional activity of this putative *Ostn* enhancer responded to *Sp7* over-expression in heterologous reporter assays (Supplemental Fig. 9c). While both SP7 cDNAs were equally expressed (Fig. 7d, right), SP7^R316C^ was defective at transactivating this *Ostn* reporter element (Fig. 7d, left).

Similar to *Sp7^OcyKO^* mice (Fig. 1b), bone biopsies from *SP7^R316C^* patients show high cortical porosity and increased bone turnover [15]. We analyzed osteocyte morphology in non-decalcified iliac crest bone biopsies from two *SP7^R316C^* patients and two age and sex-matched healthy patients. Using a novel silver staining protocol (see Methods), we observed reduced dendrite length and number in *SP7^R316C^* patients versus controls (Fig. 7e). Since *SP7^R316C^* patients can stand upright and survive into adolescence (unlike complete *Sp7*-null mice which die just after birth due to lack of osteoblasts [11]), our results suggest that this hypomorphic *SP7* mutation selectively interferes with the function of Sp7 in osteocytes.

## Discussion

Here we report a key role for the transcription factor Sp7 in osteocytogenesis, the process through which some osteoblasts embed themselves within mineralized bone matrix and become osteocytes. Sp7 plays a key role in early skeletal progenitors and their commitment to the osteoblast lineage [11]. Sp7 controls osteoblast-specific gene expression via directly binding to GC-rich promoter sequences [42] and indirect DNA binding in conjunction with Dlx family transcription factors [27] and NO66 histone demethylases [43]. However, *Sp7* expression persists throughout the osteoblast lineage [44], and our current findings demonstrate an important role for this transcription factor in osteocytogenesis. In osteocytes, Sp7 associates with a distinct distal enhancer DNA motif (Fig. 4f), suggesting that unique Sp7 modifications and/or cofactors explain the cell type specific function of this transcription factor at different stages in the osteoblast lineage.

While ‘master regulator’ transcription factors have been identified for other cell types in bone [10], factors responsible for osteocyte-specific gene expression patterns have proved elusive. Mef2 family transcription factors, which also play key roles in growth plate chondrocytes [45] and neurons [46], drive expression of the osteocyte-restricted transcript *Sost* [47–49]. However, mice lacking Mef2c in osteocytes show high bone mass but normal osteocyte morphology. Loss of Hox11 function in adult mice at the osteoblast-to-osteocyte transition causes defects in osteocyte morphology similar to those seen in our *Sp7^OcyKO^* mice [50]. The molecular mechanism through which Hox11 factors control osteocyte morphology remains unknown. Here, we demonstrate that Sp7 plays a vital role in orchestrating osteocyte differentiation and identify novel cell type-specific target genes responsible for this effect. Although a previous report suggested a role of Sp7 in osteocyte morphology [24], our study employs transcriptomic, epigenomic, and single cell profiling in order to define the molecular mechanism used by Sp7 to control osteocyte maturation, and shows the clinical relevance of this role.

Relatively few osteocyte-specific Sp7 target genes were identified by intersecting our transcriptomic and ChIP-seq data (Fig. 4g). Of these genes, *Ostn* over-expression led to dramatic rescues in Sp7 deficient phenotypes *in vitro* and *in vivo*. Osteocrin was initially identified based on its expression in osteoblasts and early embedding osteocytes [33, 34]. However, the biologic functions of osteocrin have mainly been studied in growth plate, muscle, and heart [51–55]. Of note, primate, but not rodent, neurons express *osteocrin* in an activity-dependent manner, and *osteocrin* expression in neurons of higher species may be linked to increased synaptic complexity [36]. Osteocrin is thought to enhance C-type natriuretic peptide receptor signaling by targeting NPR3, a CNP clearance receptor, for degradation. In doing so, osteocrin enhances CNP signaling via NPR2 and increases intracellular cGMP levels. While the signaling mechanisms used by osteocrin to promote osteocytogenesis remain unknown, it is interesting that cGMP has been linked to osteocyte survival downstream of semaphorin 3A signaling [56].

Defects in osteocyte dendrites in *Sp7^OcyKO^* mice is likely to be the cause, not effect, of increased osteocyte apoptosis (Fig. 1l-m). Aging and pharmacologic glucocorticoid exposure are two common conditions in which defects in osteocyte morphology and increased osteocyte apoptosis are noted. Whether osteocrin administration may rescue defects in these pathologic states remains to be determined. Our model of osteocrin administration cannot discriminate between effects of this peptide on dendrite formation versus maintenance. In *Sp7^OcyKO^* mice, it is likely that reduced inter-cellular connectivity leads to increased osteocyte apoptosis. Osteocyte apoptosis triggers RANKL-driven osteoclast recruitment [57, 58]; therefore, increased intracortical remodeling (Fig. 1c-d) observed in *Sp7^OcyKO^* mice is likely due to primary defects in osteocyte morphology.

Previous efforts to characterize non-hematopoietic cells in bone using single-cell profiling have failed to capture significant numbers of cells at the osteoblast-to-osteocyte transition [59–62]. Our approach combining Dmp1-Cre; tdTomato reporter mice with serial enzymatic digestions allowed us to identify a population of cells for single cell profiling enriched in mature osteoblasts, differentiating osteocytes, and some mature osteocytes. This population of cells is enriched in expression of genes previously linked to human fracture risk (Fig. 7a), highlight the key contribution of matrix-associated/embedded cells to human bone mass/strength. Multiple novel osteoblast/osteocyte subset markers were identified in this work (Fig. 6). Sp7-deficiency leads to an accumulation of mature osteocyte precursors and a relative paucity of mature osteocytes, an observation confirmed by complementary analyses designed to interrogate cellular differentiation stage. Future studies are needed to use the novel markers identified here to define the location and morphologic features of distinct osteocyte precursor subsets *in situ*.

Dendritic projections allow osteocytes to quickly communicate with each other via gap junctions in order to sense external cues that in turn regulate bone homeostasis [63]. While devoid of action potentials, osteocytes use calcium fluxes for electric coupling between cells to facilitate rapid response to mechanical cues [64]. As such, potential parallels between highly-connected osteocytes and neurons may emerge. Osteocyte-specific Sp7 target genes are enriched in neuronally-expressed transcripts (Supplementary Fig. 7b-c), likely due to common requirements for cytoskeletal factors in dendritic projections of osteocytes and neurons. While a great deal is known about the constituents of neuronal projections including axons, dendrites, and dendritic spines and the external cues that regulate neuronal connectivity [65], the transcriptional programs that drive axon and dendrite formation during neuronal development and adult neurogenesis remain less well understood [66]. Whether the Sp7-dependent gene module identified here in osteocytes also participates in neuronal morphogenesis remains to be determined. Shared patterns of gene expression between osteocytes and neurons are likely to reflect common use of fundamental cytoskeletal factors to maintain cellular projections. It is possible that the Sp7-dependent signature identified here also participates in development and maintenance of cellular projections in ‘dendrite’-bearing cell types in other organs [67].

Taken together, our findings demonstrate that Sp7, a gene linked to rare and common human skeletal traits, orchestrates osteocytogenesis via a suite of target genes that promote the optimal formation and maintenance of osteocyte dendrites. Osteocrin, a secreted factor positively regulated by Sp7, can rescue osteocyte morphology and survival defects in Sp7-deficient mice. These findings highlight shared features between osteocytic and neuronal connectivity and highlight novel steps in osteocytogenesis that may be targeted to improve bone strength for individuals with osteoporosis.

---

**Fig. S1:**
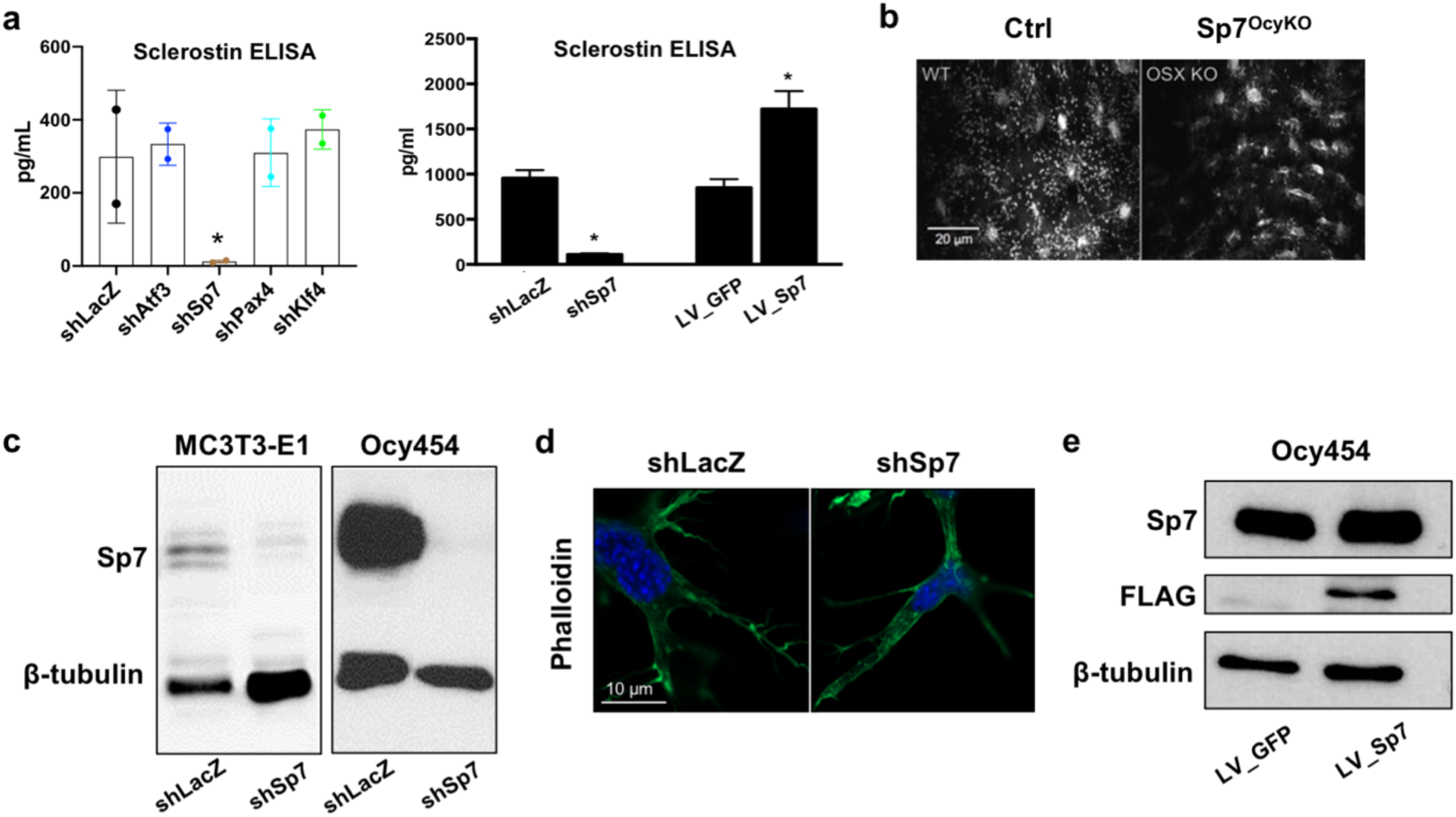
(**a**) Left panel: Ocy454 cells were infected with shRNA-expressing lentiviruses targeting the indicated gene followed by puromycin selection. Cells were grown at 37°C for 14 days, then sclerostin ELISA was performed on 72h conditioned medium. Only *Sp7* knockdown reduced sclerostin level. Right panel: Ocy454 cells were subjected to *Sp7* knockdown (left) or *Sp7* overexpression (right) followed by sclerostin ELISA. (**b**) Representative *in vivo* third harmonic generation images from control and *Sp7^OcyKO^* mice, revealing reduced punctate signal representing osteocyte dendrites. See also Fig. 1f. (**c**) MC3T3-E1 and Ocy454 cells were infected with the shRNA-expressing lentiviruses followed by immunoblotting as indicated. shSp7 infection leads to near complete loss of Sp7 protein. (**d**) Ocy454 cells were infected with control or *Sp7* knockdown lentiviruses followed by growth in 3D culture. Similar to results seen in MC3T3-E1 cells (**c**), *Sp7* knockdown causes defective formation of phalloidin-positive long filaments in this cell type. Scale bar represents 10 µm. (**e**) Ocy454 cells were infected with control (LV-GFP) or FLAG-Sp7 expressing lentiviruses, followed by immunoblotting as indicated.

**Fig. S2:**
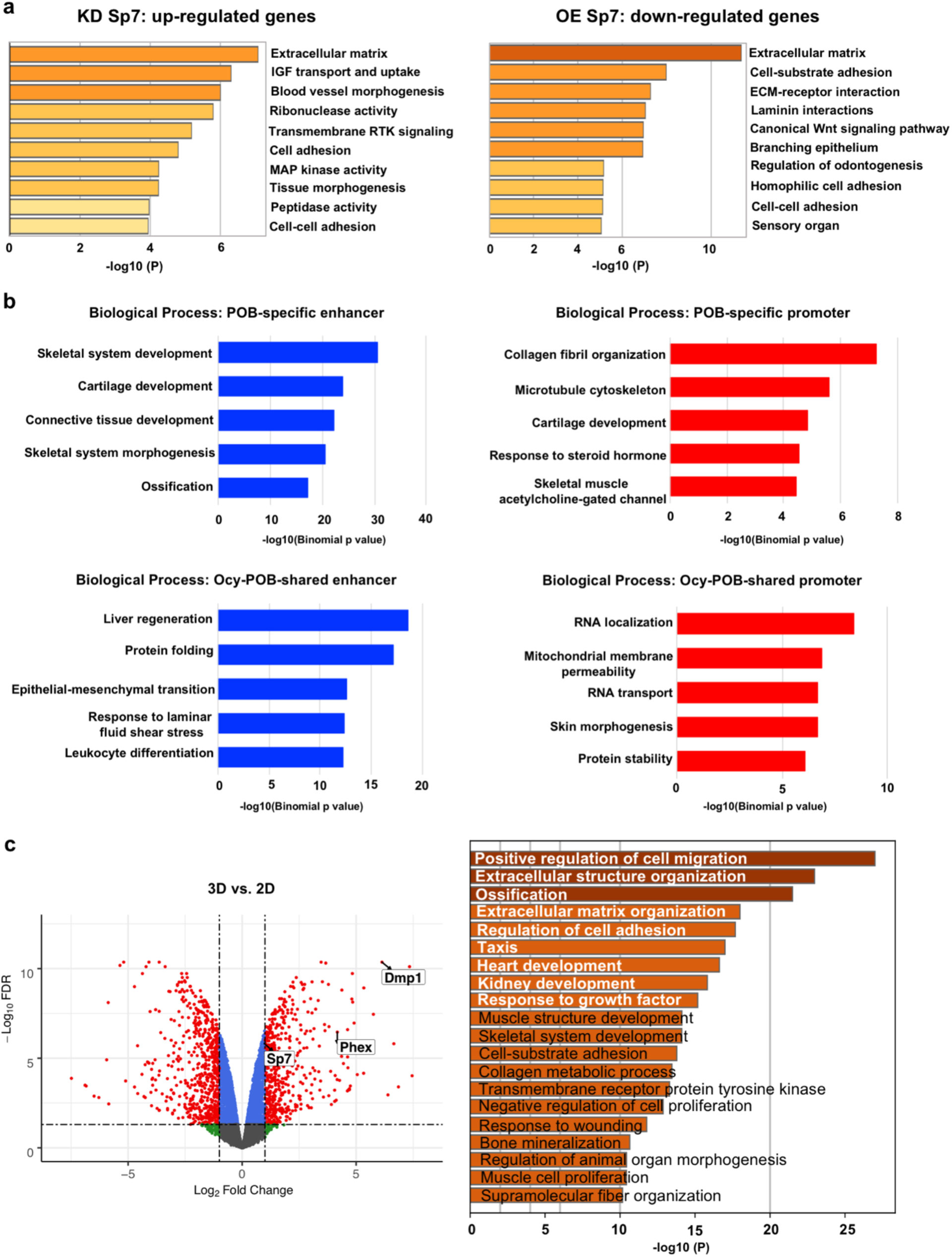
(**a**) Gene ontology analysis of genes up-regulated in response to *Sp7* knockdown (KD, left) and down-regulated in response to *Sp7* overexpression (OE, right). (**b**) Top: functional analysis of genes bound by Sp7 only in osteoblasts (POB) at the enhancer (left) and promoter (right) region. Bottom: functional analysis of genes bound by Sp7 in both osteoblasts and Ocy454 cells at the enhancer (left) and promoter (right) regions. (**c**) Left: volcano plot from bulk RNA-seq data of MC3T3-E1 cells infected with shLacZ. Cells were grown in either 2D or 3D culture. Differentially expressed genes are defined as log2 FC >1 or <-1 with adjusted p value (FDR) < 0.05. Several osteocyte markers are highlighted: *Sp7*, *Dmp1* and *Phex*; Right: gene ontology analysis of differentially expressed genes in 3D culture reveals enrichment in several terms associated with cell migration, ossification and supramolecular fiber organization.

**Fig. S3:**
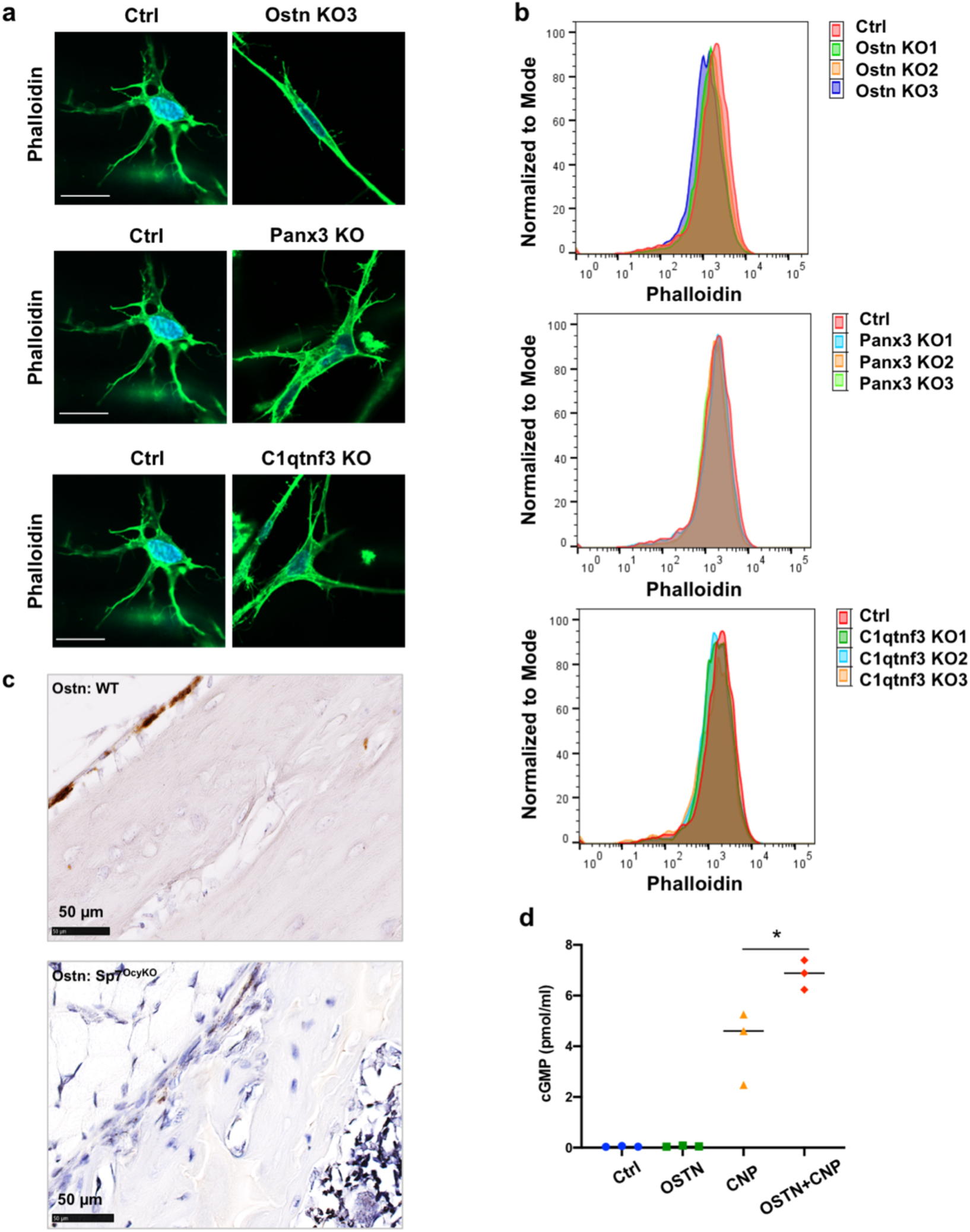
(**a,-b**) Ocy454 cells were subjected to CRISPR/Cas9-mediated deletion of the indicated gene, followed by growth in 3D culture conditions and phalloidin staining (**a**) or flow cytometry (**b**). Only *Ostn* CRISPR deletion led to defects in F-actin content. Scale bars represent 10 µm. (**c**) RNA *in situ* hybridization of *Ostn* in 6-week-old WT (top) and Sp7^OcyKO^ (bottom) mouse tibia. (**d**) cGMP levels in Ocy454 cells were measured by ELISA. Cells were incubated for 30 minutes under control conditions (HBSS) or with 50 nM CNP, or 500 nM OSTN, or 50 nM CNP + 500 nM OSTN. Data are representative of 3 independent experiments.

**Fig. S4:**
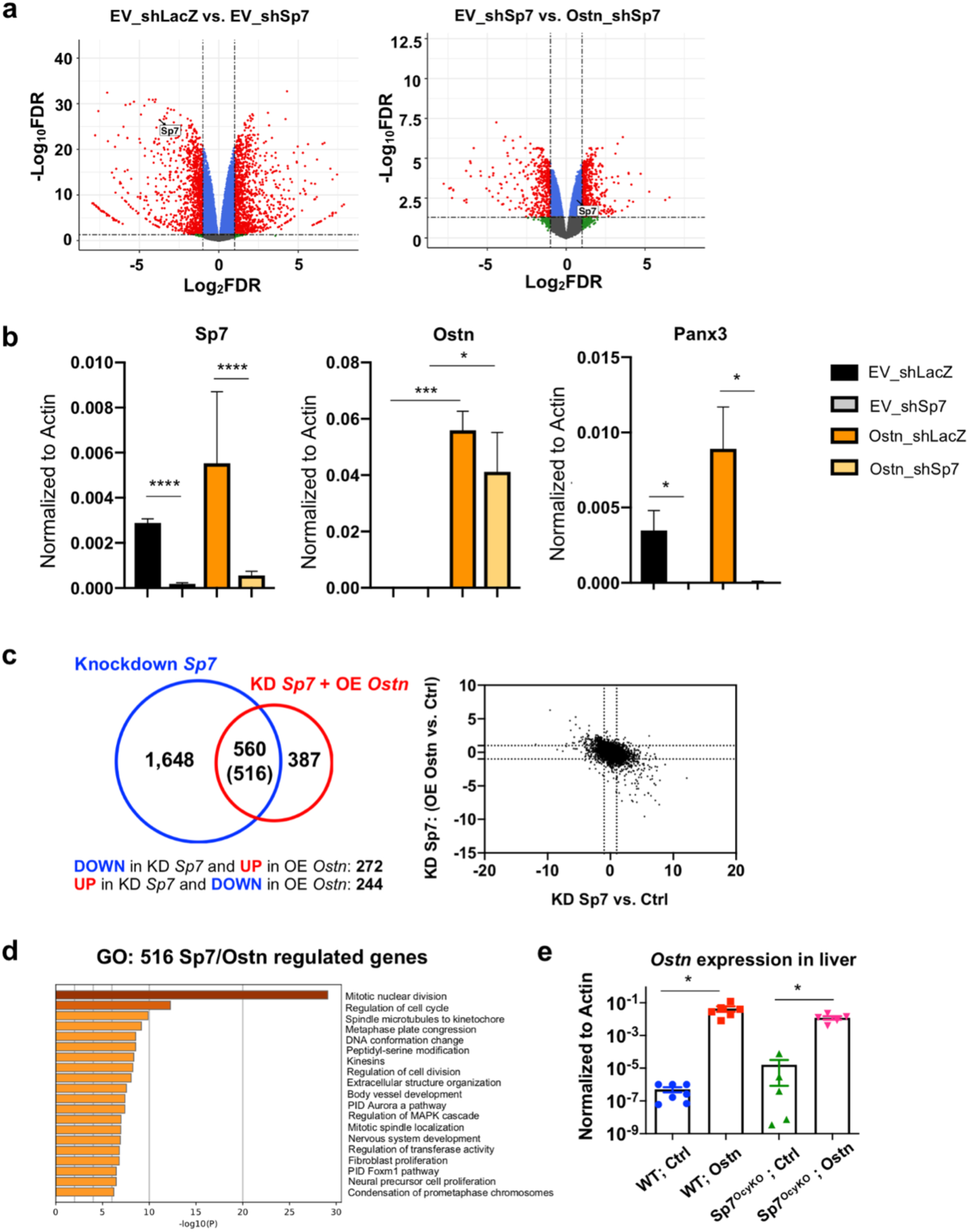
(**a**) Left: volcano plot from bulk RNA-seq data of MC3T3-E1 cells infected with shSp7 versus control (shLacZ). Right: volcano plot from bulk RNA-seq data of MC3T3-E1 cells infected with shSp7 then subjected to control (EV, empty vector) or *osteocrin* (*Ostn*) over-expression. Differentially expressed genes are defined as log2 FC >1 or <-1 with adjusted p value (FDR) < 0.05. (**b**) Cells as in (**a**) were analyzed by RT-qPCR for the indicated genes, demonstrating expected degrees of *Sp7* knockdown and *Ostn* over-expression. *Panx3* is a Sp7-dependent gene whose expression is not rescued by Ostn. (**c**) Left, RNA-seq was performed in control (shLacZ) versus *Sp7* knockdown MC3T3-E1 cells (blue circle), and *Sp7* knockdown cells rescued with control (LV-empty vector) or osteocrin cDNA (red circle). The intersection of differentially-expressed genes in these two comparisons revealed 560 common transcripts, the majority (516) of which were discordantly regulated by the two perturbation. Right, scatterplot analysis between the effects of *Sp7* knockdown and *osteocrin* expression in *Sp7* knockdown cells. (**d**) Gene ontology analysis of 516 genes regulated in opposite directions by *Sp7* knockdown and *Ostn* overexpression in *Sp7* knockdown cells. GO enrichment analysis revealed that these counter-regulated genes are strongly involved in cell cycle and microtubule cytoskeleton organization. This indicates that there is only a small portion of Sp7/Ostn downstream genes that function in osteocyte dendrite formation. See also Supplementary Table 6. (**e**) RNA was isolated from liver of mice infected with control AAV8 or AAV8-mOstn. Hepatic *Ostn* expression is very low in control mice, and robustly induced after AAV8-mOstn infection (note log10 y-axis scale).

**Fig. S5:**
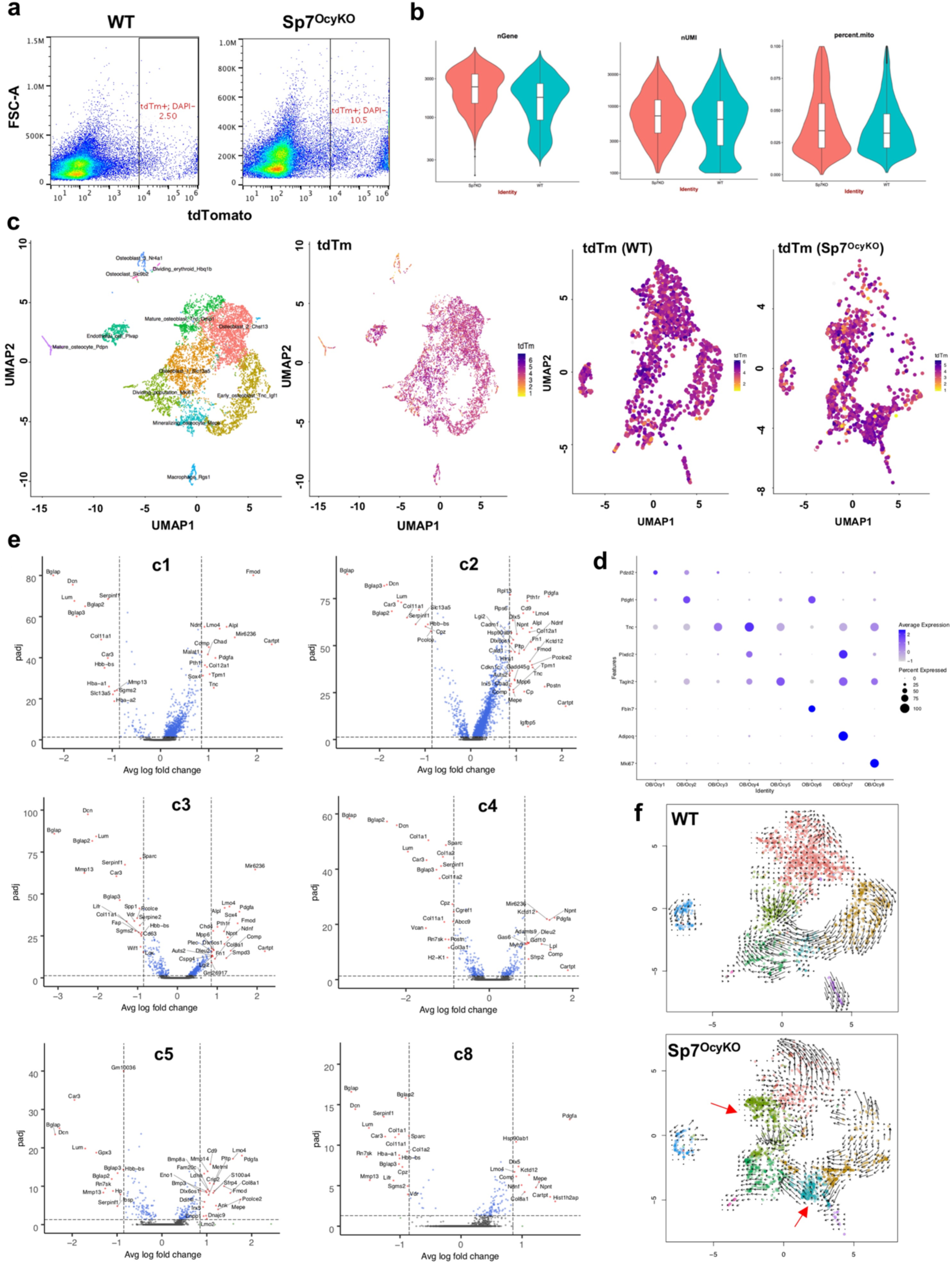
(**a**) Flow cytometry plots showing tdTomato expression in DAPI-negative cells obtained from fractions 5, 7, and 8 after serial collagenase/EDTA digestions and then sorted for single-cell RNA-sequencing. (**b**) Quality control metrics from WT and *Sp7^OcyKO^* libraries showing acceptable distribution of genes detected per cell, unique molecular identifiers per cell, and mitochondrial read abundance. (**c**) First panel (from left to right): clustering of all cells sequenced reveals main populations of osteoblast-lineage cells, along with other cell types including endothelial cells, macrophages, osteoclasts, and dividing erythroid cells. Second panel: tdTomato expression was mapped to all cells sequenced. Of note, some non-osteoblast lineage cells (endothelial cells, macrophages) do express detectable tdTomato mRNA. These clusters plus osteoclasts and dividing erythroid cells were excluded from future analyses. Third and fourth panels: tdTomato expression in remaining osteoblast-lineage cells shows comparable expression levels of this Dmp1-Cre-dependent reporter in WT and *Sp7^OcyKO^* cells. (**d**) Dotplot showing the expression of top1 marker in each cluster. (**e**) Cluster-specific differential gene expression volcano plots for indicated clusters. The x-axis shows log2 fold change (*Sp7* mutant versus WT) and the y-axis shows the adjusted p value. (**f**) Velocyto trajectory analysis in control and *Sp7^OcyKO^* libraries shows lack of orderly progression to the terminal osteocyte stage (cluster 6 in purple) in *Sp7* mutant mice. Red arrows again indicate apparent differentiation arrest in c3 and c5. See also Fig. 6g.

**Fig. S6:**
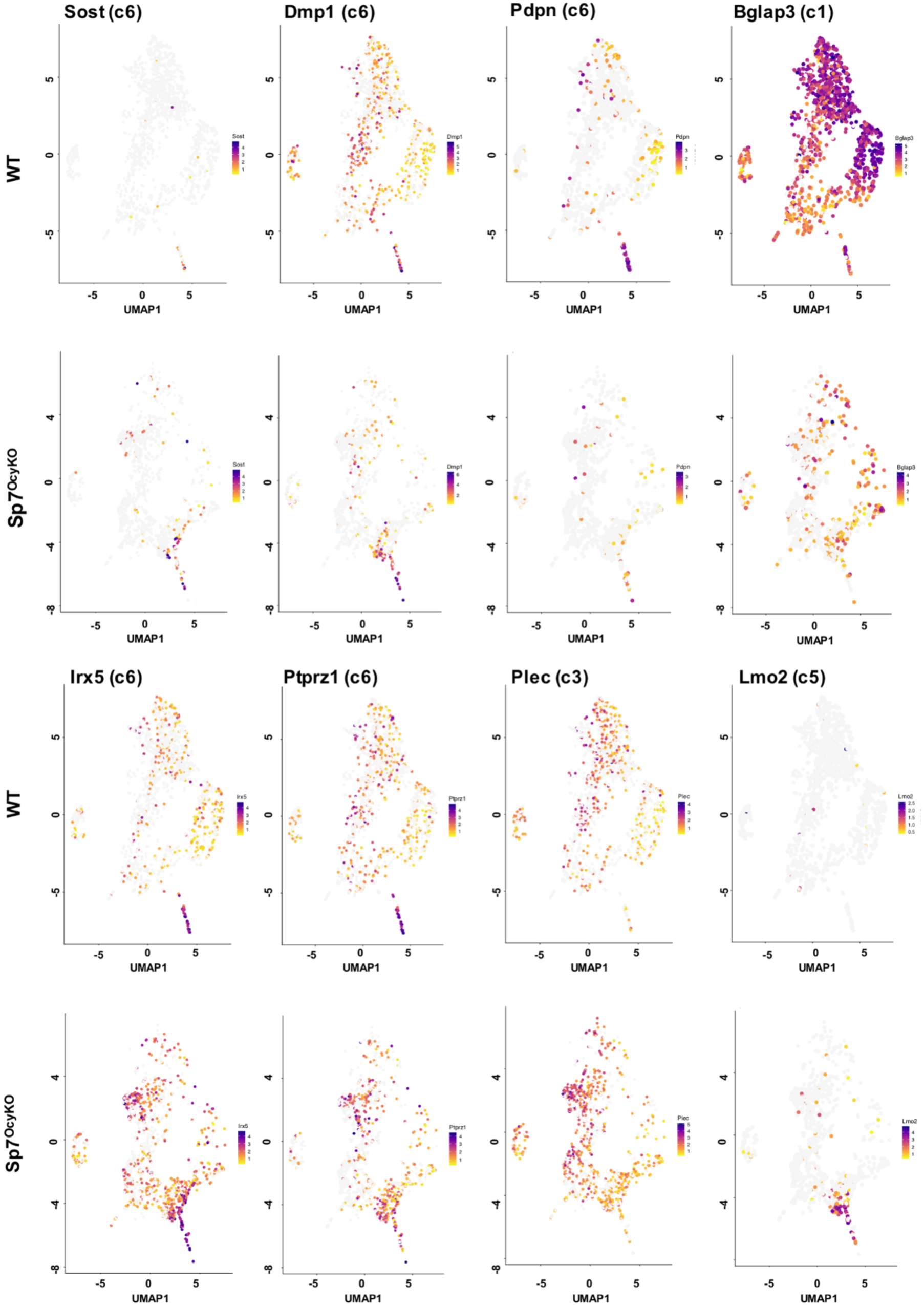
Feature plots showing relative transcript abundance of indicated genes. Top: expression of canonical osteocyte (*Sost, Dmp1, Pdpn*) and osteoblast-lineage (*Bglap3*) marker genes. As expected, *Bglap3* is highly expressed in all cells analyzed. In WT libraries, osteocyte marker expression is restricted to cluster 6, while dysregulated expression of *Sost*, *Irx5*, and *Ptprz1* is noted in *Sp7* mutant libraries, especially in *Lmo2*-expressing cells in cluster 5 and *Plec*-expressing cells in cluster 3. In contrast, other terminal osteocyte markers (*Pdpn* and *Ptprz1*) are restricted to osteocytes in control mice and essentially absent in *Sp7* mutants.

**Fig. S7:**
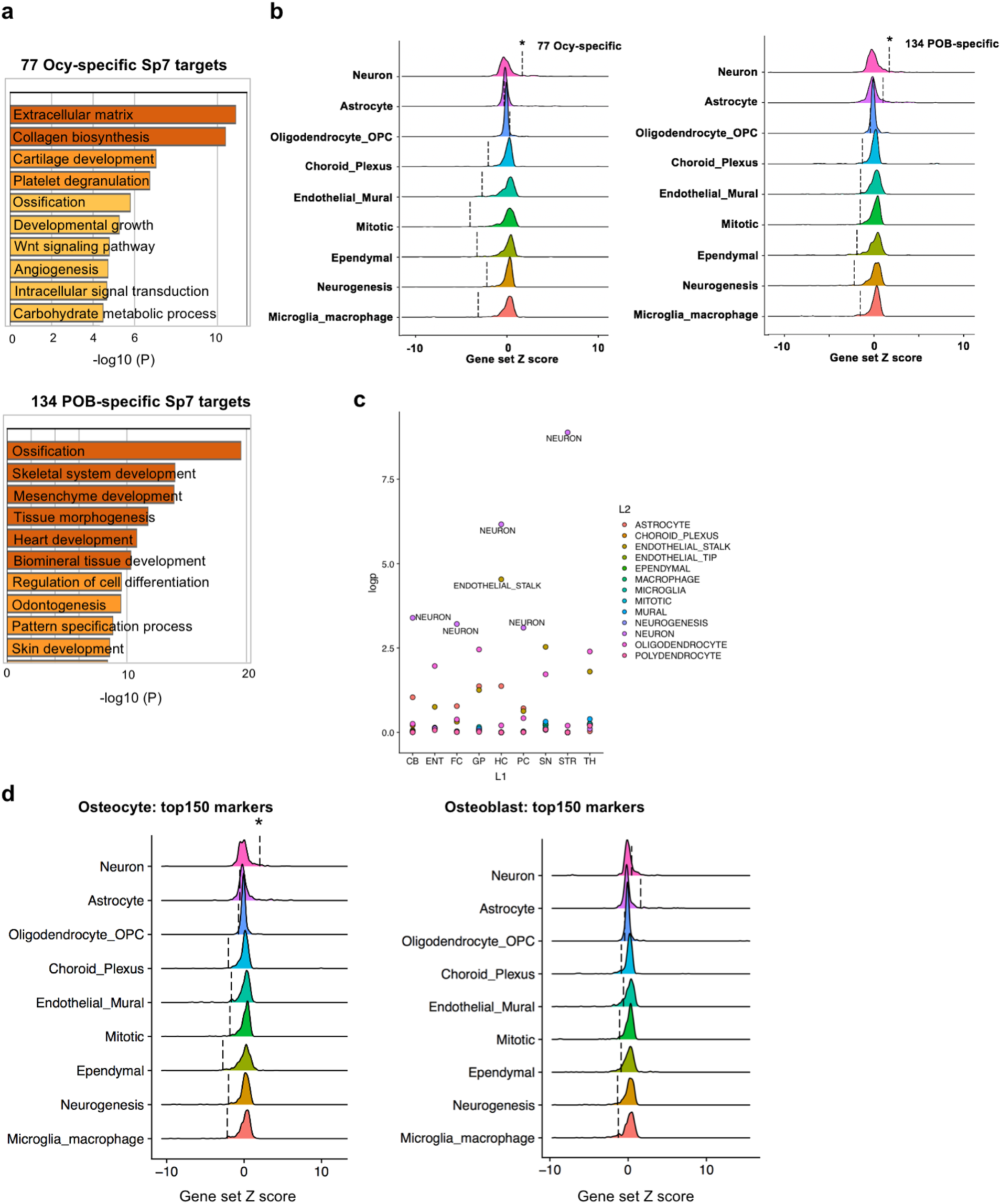
(**a**) GO enrichment of 77 osteocyte-specific (top) and 134 primary osteoblast-specific (bottom) Sp7 targets. (**b**) Left: The expression of 77 osteocyte-specific Sp7 target genes was analyzed in a mouse brain single-cell RNA-seq atlas, where significant enrichment is found in aggregated expression values for neurons (p = 0.047) as compared against all other major cell type aggregated expression values. OPC: Oligodendrocyte progenitor cell; Mitotic: Mitotic cells, Neurogenesis: Neurogenesis-associated cells. Right: 134 genes (Supplementary Table 5) are revealed by intersecting osteoblast-specific genes derived from primary osteoblasts (FDR < 0.05, 951 genes) and genes with associated primary osteoblast specific Sp7 ChIP-seq peaks (1,962 genes). The expression of this set of 134 genes was analyzed in a mouse brain single-cell RNA-seq atlas and was significantly enriched in neurons (p = 0.045) compared other brain cell types. (**c**) Enrichment score of 77 gene set expression in cell types across nine mouse brain regions. CB - cerebellum, ENT - entorhinal cortex, FC - frontal cortex, GP - globus pallidus, HC - hippocampus, PC -parietal cortex, SN - substantia nigra, STR - striatum, TH - thalamus. Across multiple brain regions, neurons show relative enrichment of this gene set versus other non-neuronal cell types. (**d**) Left: the expression of top150 mature osteocyte markers (c6) was analyzed in a mouse brain single-cell RNA-seq atlas, where significant enrichment is found in aggregated expression values for neurons (p = 0.022) as compared against all other major cell type aggregated expression values. Right: the expression of top150 osteoblast marker (c1+2) was analyzed in a mouse brain single-cell RNA-seq atlas. OPC: Oligodendrocyte progenitor cell; Mitotic: Mitotic cells, Neurogenesis: Neurogenesis-associated cells.

**Fig. S8:**
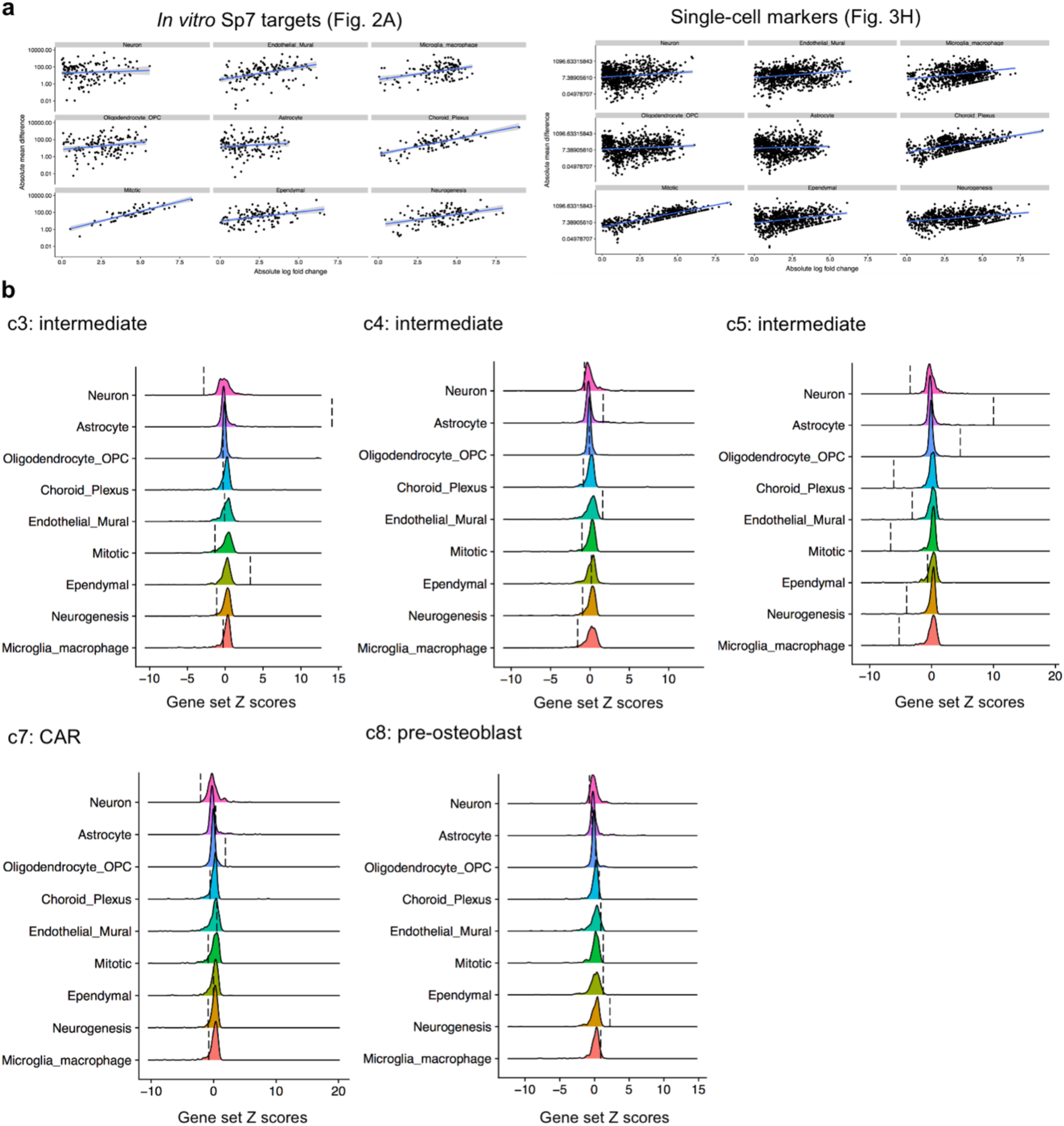
(**a**) Correlation between brain cell type mean difference and log_2_FC of *in vitro Sp7* targets (left, Supplementary Figure 7b) and top150 single-cell RNA-seq markers (right, Supplementary Figure 7d). See Methods for details. (**b**) The expression of top150 markers from other osteo-lineage clusters (c2-c5, c7-8) was analyzed in a mouse brain single-cell RNA-seq atlas. None of them show significant enrichment in neurons compared to other brain cell types. OPC: Oligodendrocyte progenitor cell; Mitotic: Mitotic cells, Neurogenesis: Neurogenesis-associated cells.

**Fig. S9:**
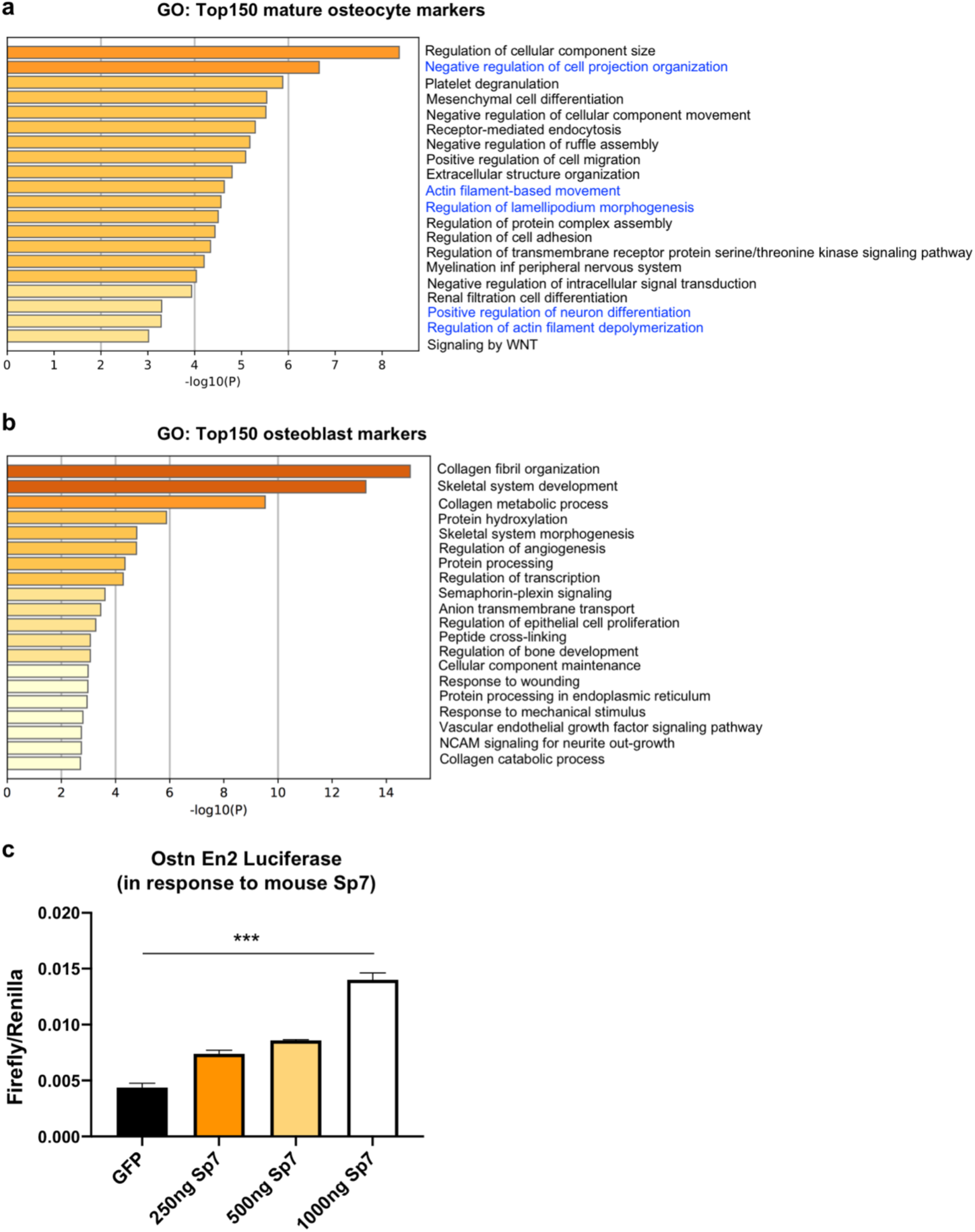
(**a-b**) GO enrichment terms of top150 mature osteocyte markers derived from cluster 6 and top150 canonical osteoblast markers derived from cluster 1+2. (**c**) Luciferase assay in 293T cells showing effects of murine *Sp7* overexpression on *osteocrin* enhancer 2 activity. ***, p<0.001.

## Acknowledgements

We thank Drs. Tatsuya Kobayashi, Amar Sahay, Jay Rajagopal, Lauren Surface, Ryan Logan, Marianne Seney, and all members of the Wein laboratory for discussions. We thank Dr. Benoit de Crombrugghe for providing *Sp7* conditional knockout mice, Dr. Vanda Jorgeti for providing healthy age-matched control iliac crest samples, and Dr. Frank Rauch for providing iliac crest biopsy samples from *Sp7^R316C^* mutant patients. MNW acknowledges funding support from the MGH Department of Medicine (Transformative Scholars award), the American Society of Bone and Mineral Research (Rising Star Award), and the National Institute of Health (AR067285). µCT and bone histomorphometry were performed by the Center for Skeletal Research, an NIH-funded program (P30 AR066261). Confocal microscopy was supported by the NIH Shared Instrumentation Grant (SIG) S10OD021577.

## Author contributions

JSW, FM, DR, HH, CDC, RP, NG, TE, MB, DJB, DT, AA, and MNW conceived study design and performed experiments. JSW, TK, FM, HH, EZM, DT, CPL, HMK, and MNW analyzed data. MF, CFM, and MF contributed key reagents. JSW and MNW wrote the manuscript. All authors edited and approved the manuscript.

## Resource availability

### Lead contact

Further information and requests for resources and reagents should be directed to and will be fulfilled by the Lead Contact, Marc Wein.

### Materials availability

Unique materials and reagents generated in this study are available upon request from the Lead Contact.

### Data and code availability

The RNA-seq, ChIP-seq and scRNA-seq data will be deposited in NCBI’s Gene Expression Omnibus (GEO) (GSE154719). The authors declare that all other data supporting the findings of this study are available within the article and its supplementary information files.

## Methods

### Mice

*Dmp1*-*Cre* (9.6 kB) transgenic mice ([22]; RRID: MGI:3784520) and Ai14 Cre-dependent tdTomato reporter (JAX, catalog number 007914) were intercrossed. Floxed *Sp7* mice were kindly provided by Dr. Benoit de Crombrugghe [68]. *Dmp1-Cre; tdTm^+^* mice were crossed with *Sp7^fl/+^* mice to generate *Sp7^OcyKO^* (*Dmp1-Cre; Sp7^fl/fl^; tdTm^+^*) and control (*Dmp1-Cre; Sp7^+/+^; tdTm^+^*) mice. Genotypes were determined by PCR using primers listed in Supplementary Table 8. All mouse strains were backcrossed to C57BL/6J for at least 4 generations. While this degree of backcrossing may be insufficient for subtle skeletal phenotypes, the phenotype of *Sp7^OcyKO^* is quite striking. Moreover, littermate controls were used for all studies. Both males and females were included in this study. All procedures involving animals were performed in accordance with guidelines issued by the Institutional Animal Care and Use Committees (IACUC) in the Center for Comparative Medicine at the Massachusetts General Hospital and Harvard Medical School under approved Animal Use Protocols (2019N000201). All animals were housed in the Center for Comparative Medicine at the Massachusetts General Hospital (21.9 ± 0.8 °C, 45 ± 15% humidity, and 12-h light cycle 7 am–7 pm).

### Cell culture

Cells were passed in alpha-MEM supplemented with heat inactivated 10% fetal bovine serum and 1% antibiotic-antimycotic (Gibco^TM^) with 5% CO_2_. Ocy454 cells [21] were plated at 10^5^ cells ml^−1^ to reach confluence at the permissive temperature (33 °C) in 2-3 days. Ocy454 cells express a thermosensitive large T antigen which is active at 33 °C and inactive at 37 °C. Subsequently, cells were switched to the non-permissive temperature (37 °C) to promote osteocyte differentiation. For protein and gene expression analyses, cells were analyzed after culture at 37 °C for 14-21 days. For microscopy and flow cytometry analyses, cells were cultured at 37 °C for 5-7 days prior of staining. MC3T3-E1 cells (subclone 4, ATCC CRL-2593) were plated at 10^5^ cells ml^−1^ to reach confluence and were kept at 37 °C for the indicated time points.

For 3D culture, rat tail type I collagen (Advanced BioMatrix, 5153) was first mixed with the neutralized buffer on ice (neutralized buffer: 1.7 ml of 3M NaOH, 10 ml of 1 M HEPES, 1.1 g of NaHCO_3_, up to 50 ml of dH_2_O) to reach pH = 7. Cells were diluted to 5 x 10^5^ cells ml^−1^ before mixing with the collagen-buffer mix on ice (1:1 volume). The cell/collagen/buffer mixture was plated in 12-well or 24-well plates and incubated in 37 °C for 10 minutes to polymerize. Equal volume of culture media (alpha-MEM supplemented with heat inactivated 10% fetal bovine serum and 1% antibiotic-antimycotic, Gibco^TM^) was added on top of the solidified gel prior to subsequent culture.

### shRNA infection and lentiviral transduction

See Supplementary Table 8 for all shRNA sequences used. For shRNA, lentiviruses were produced in HEK293T cells in a pLKO.1-puro (Addgene, plasmid 8453) backbone. Viral packaging was performed in 293T cells using standard protocols (http://www.broadinstitute.org/rnai/public/resources/protocols). Briefly, HEK293T cells were plated at 2.2 x 10^5^ ml^−1^ and transfected the following day with shRNA-expressing plasmid (shSp7 or shLacZ plasmid) along with psPAX2 (Addgene, plasmid 12260) and pMD2.G (Addgene, plasmid 12259) using Fugene-HD (Promega). Medium was changed the next day and collected 48 hours later. Ocy454 cells were exposed to lentiviral particles overnight at 33°C in the presence of polybrene (2.5 µg ml^−1^). Media was then changed with puromycin (4 µg ml^−1^). MC3T3-E1 cells were exposed to lentivirus overnight at 37 °C. Cells were maintained in selection medium throughout the duration of the experiment.

Mouse FLAG-Sp7 or osteocrin cDNAs were introduced via lentivirus via pLX_311 backbone (Addgene, plasmid 118018) as previously described [21]. Briefly, mouse Sp7 and GFP lentiviruses were generated by transfecting HEK293T cells with a blasticidin resistance backbone (Addgene, plasmid 26655) along with psPAX2 and MD2.G. Ocy454 and MC3T3-E1 cells were infected with lentiviral particles expressing GFP or Sp7. After 24 hours, cells were selected with blasticidin (4 µg ml^−1^) and used for subsequent experiments. To overexpress *Ostn* in *Sp7* knockdown cells, *Sp7* knockdown cells were exposed to Ostn or EV lentiviral particles overnight at 37 °C in the presence of polybrene (2.5 µg ml^−1^). Media was changed with puromycin (4 µg ml^−1^) and blasticidin (4 µg ml^−1^) on the next day. FLAG-hSP7, FLAG-hSP7^R316C^ and EF1a-eGFP constructs were synthesized *de novo* (VectorBuilder).

### Adeno-associated virus 8 (AAV8) infections

AAV8 vectors encoding codon mouse *Ostn* or green fluorescent protein (GFP) under the control of the chicken beta actin (CAG) promoter (AAV8-CAG-mOstn-WPRE and AAV8-CAG-eGFP vectors) were generated (Vector Biolabs). 3-week-old mice were injected with AAV8 by intraperitoneal (IP) injection at a dose of 5 × 10^11^ gc per mice in a total volume of 100 µl.

### Human bone sample silver impregnation and osteocyte histomorphometry

Control bone samples were obtained from the bone biopsies bank of the LIM 16-Laboratório de Fisiopatologia Renal, Hospital das Clínicas HCFMUSP, Faculdade de Medicina-Universidade de São Paulo. Transiliac bone biopsies were performed 3-5 days after a course of double labeling with tetracycline (20 mg kg^−1^ per day for 3 days) with a 10-day interval. Undecalcified specimens were fixed in 70% ethanol, dehydrated, embedded in methyl methacrylate and cut using a tungsten carbide knife. 5 μm-thick sections were stained with Goldner’s Trichrome for static bone parameters. Silver nitrate impregnation was used for osteocytes lacunae and its dendrites appreciation. The sections were deacrylated in cold MMA for 10-15 minutes. After rehydration in graded alcohol solutions, 5 μm-thick sections were decalcified in a 20% EDTA (pH = 7.8) for 30 minutes and incubated with 200 mM silver nitrate in solution containing 0.6% gelatin type A and 0.3% of formic acid for 30 minutes. The sections were then washed and developed in an aqueous 15% solution of sodium thiosulfate (Na_2_S_2_O_3_) for 10 minutes. Unstained 10-μm-thick sections were analyzed under UV light for dynamic parameters. Quantification of the dendrite number and length were performed with ImageJ by thresholding gray-scale images for dark, silver-stained lacunae and canaliculi. All histomorphometric analyses were performed using a semi-automatic image analyzer and OsteoMeasure software (OsteoMetrics, Inc., Atlanta, GA, USA), at 125x magnification, and the full bone structure located between the two cortical areas was evaluated.

### Histology and immunohistochemistry

Formalin-fixed paraffin-embedded decalcified tibia sections from 6-week-old and 8-week-old mice were obtained. For anti-Sp7 immunohistochemistry staining, antigen retrieval was performed using proteinase K (20 μg ml^−1^) for 15 minutes. Endogenous peroxidases were quenched, and slides were blocked in TNB buffer (PerkinElmer), then stained with anti-Sp7 antibody (Abcam, ab22552) at a concentration of 1:200 for 1 hour at room temperature. For activated caspase-3 IHCs, sections were stained with primary antibody (Cell Signaling Technology, 12692) at 1:500 overnight at 4 °C. Sections were washed, incubated with HRP-coupled secondary antibodies, signals amplified using tyramide signal amplification and HRP detection was performed using 3,3′-diaminobenzidine (DAB, Vector Laboratories) for 2–3 minutes. Slides were briefly counterstained with hematoxylin before mounting. Hematoxylin and eosin (H&E) staining was performed using standard protocols. Quantification of Sp7 positive osteocytes and empty lacunae were performed in ImageJ imaging software on the blind-test manner.

### Immunoblotting and ELISA

Immunoblotting was performed as previously described [69, 70]. For both Ocy454 and MC3T3-E1 *Sp7* knockdown and overexpressing cells, whole cell lysates were prepared using TNT (Tris-NaCl-Tween buffer, 20 mM Tris-HCl pH = 8, 200 mM NaCl, 0.5% Triton X-100 containing protease inhibitor (PI), 1 mM NaF, 1 mM DTT, 1 mM vanadate). Adherent cells were washed with ice cold PBS, then scraped into TNT buffer on ice. Material was then transferred into Eppendorf tubes kept on ice, vortexed at top speed for 30 seconds, then centrifuged at top speed for 6 minutes at 4 °C. For immunoblotting, lysates were separated by SDS-PAGE and proteins were transferred to nitrocellulose. Membranes were blocked with 5% milk in tris-buffered saline plus 0.05% Tween-20 (TBST) and incubated with primary antibody overnight at 4°C diluted in TBST plus 5% BSA. The next day, membranes were washed, incubated with appropriate HRP-coupled secondary antibodies (Anti-rabbit HRP, Cell Signaling Technology 7074, 1:2000), and signals detected with ECL Western Blotting Substrate (Pierce) or ECL Plus Western Blotting Substrate (Pierce). The primary antibodies were Sp7 (Abcam, ab22552), DYKDDDDK tag (Cell Signaling Technology, 2368), GAPDH (Cell Signaling Technology, 2118) and β-tubulin (Cell Signaling Technology, 5346). For transfected HEK293T cells, protein lysates were collected 48 hours after transfection as above.

### Sclerostin ELISA

Sclerostin ELISAs were performed using an antibody pair (Scl Ab-VI and Scl Ab-VII) as described previously [21]. Conditioned medium (48-72 hours) was harvested from Ocy454 cells as indicated in the figure legends and stored at −80 °C until further use. High binding 96 well plates (Fisher, 21-377-203) were coated with Scl Ab-VI capture antibody (3 µg ml^−1^) in PBS for one hour at room temperature. Plates were washed (PBS plus 0.5% Tween-20) and blocked with wash buffer supplemented with 1% BSA and 1% normal goat serum for one hour at room temperature. Samples (60 µl well^−1^) were then added along with a standard curve of murine recombinant Sclerostin (Alpco, Salem, NH) and plates were incubated overnight at 4 °C. Plates were washed three times, and then incubated with HRP-coupled Scl Ab-VII detection antibody (0.5 µg ml^−1^) for one hour at room temperature. After washing, signal detection was performed using Ultra TMB ELISA (Pierce, Rockford, IL), stopped by 2N sulfuric acid, and read at 450 nm. Prior to harvesting supernatant, cell number per well was always determined using PrestoBlue assay (Life Technology) read at 570 nm and 600 nm according to the manufacturer’s instructions.

### Measurements of intracellular cGMP

Ocy454 cells, grown in 96-well plates, were incubated at 37 °C for 2 days. Cells were subsequently stimulated with incubation medium (Hank’s Balanced Salt solution with 0.075% BSA, 10 mM HEPES and 2 mM IBMX) containing 50 nM human CNP or 500 nM human OSTN at room temperature for 30 minutes. Cells were collected in 0.1 M HCl. The amount of cGMP in each cell lysate was measured using the Cyclic GMP ELISA Kit (Cayman Chemical) according to the manufacturer’s protocol.

### Histomorphometry

Right tibiae from 8-week-old mice were subjected to bone histomorphometric analysis (calcein labeling). All mice received intraperitoneal calcein injections (Sigma Aldrich, 20 mg kg^−1^) at 2 days prior to sacrifice. The tibia was dissected and fixed in 70% ethanol for 3 days. Fixed bones were dehydrated in graded ethanol, then infiltrated and embedded in methylmethacrylate without demineralization. Undecalcified 5 μm longitudinal sections were obtained using a microtome (Leica Biosystems, RM2255). The observer was blinded to the experimental genotype at the time of measurement. Digital images were obtained via fluorescent microscopy.

### Silver staining of mouse osteocyte morphology

As described previously [71], paraffin-embedded mouse tibia sections were deparaffinized and incubated in two parts 50% silver nitrate and one part 1% formic acid in 2% gelatin solution for 55 minutes. Stained slides were then washed in 5% sodium thiosulfate for 10 minutes and subsequently dehydrated, cleared, and mounted. Quantification of the dendrite number and length were performed with ImageJ by thresholding gray-scale images for dark, silver-stained lacunae and canaliculi.

### TUNEL staining

Formalin-fixed paraffin-embedded decalcified tibia sections from 6-week-old and 8-week-old mice were obtained. For Terminal deoxynucleotidyl transferase dUTP nick end labeling (TUNEL), sections were fixed with 4% PFA (Pierce^TM^, 28908) in PBS for 15 minutes at room temperature and permeabilized with PCR grade recombinant Proteinase K (Roche Applied Science, 3115887001) for 30 minutes at 37 °C. Apoptotic cells were examined with Roche *in situ* cell death detection kit (Roche Applied Science, 11684795910) according to instructions and followed by DAPI (Invitrogen) for fluorescent microscopy.

### RNAscope

Formalin-fixed paraffin-embedded decalcified tibia sections (5 μm) from 6-week-old mice were obtained. Bone sections were processed for RNA in situ detection using RNAscope 2.l5 HD Assay-Brown (Chromogenic) according to the manufacturer’s instructions (Advanced Cell Diagnostics, [72]). For antigen retrieval, bone sections were pretreated with hydrogen peroxide and pepsin (1 hour at 40 °C, Sigma-Aldrich). Tissue sections were hybridized with target probes (2 hours at 40 °C), amplified (Amp1-4: 40 °C; Amp5-6: room temperature) and chromogenic detected using DAB followed by counterstaining with hematoxylin (American Master Tech Scientific). RNAscope probes used were: *Ostn* (NM_198112.2, region 2-1144), *Tnc* (NM_011607.3, region 875-1830), *Kcnk2* (NM_001159850.1, region 734-1642), *Dpysl3* (NM_001291455.1, region 772-1884), *Fbln7* (NM_024237.4, region 198-1595) and *Dabp* (negative control). Representative figures are from n=3 wild-type C57B6 mice.

### Micro-CT

Femurs were harvested from 6-week and 8-week old mice after fixed with 10% formalin for one day and stored in 70% ethanol. Micro-CT imaging was performed on a bench-top scanner (µCT 40, Scanco Medical AG, Brüttisellen, Switzerland) to measure the morphology of the femoral mid-diaphysis (Scanning parameters: 10μm isotropic voxel, 70 kVp, 114 mA, 200 ms integration time). A 500μm long region of the mid-diaphysis was scanned, bone segmented from surrounding tissue using a threshold of 700 mgHA cm^−3^ and the geometry of the cortex analyzed using the Scanco Evaluation Program.

### Third harmonic generation imaging

To image bone/interstitial fluid boundaries surrounding osteocytes in calvariae, *Sp7^OcyKO^* and control mice were anesthetized with isoflurane, and placed in a custom 3D printed heated mouse holder. An incision was made on the top of the skull. Skin was partly detached exposing the periosteum and the calvarial bone [23]. A microscope cover glass holder was used to hold a microscope cover glass over the dissected area, which was hydrated with PBS. The imaging locations were consistently chosen in regions 3, 4, 5 and 6 [73], 100–300 μm away from the coronal and central veins, while imaging depths were chosen, starting from ∼10 μm below the bone surface. A turn-key laser with a 5 MHz repetition rate, 370 fs duration pulses and 1550 nm wavelength (FLCPA-01CCNL41, Calmar) was coupled to a 60 cm polarization maintaining large mode area single mode photonic crystal fiber with a 40 μm diameter core (LMA40, NKT Photonics, Birkerød, Denmark) which was coupled to an in-house laser scanning microscope. The microscope consisted of a polygonal laser scanner (DT-36-290-025, Lincoln Laser, Phoenix, AZ) for the fast axis and a galvanometric scanning mirror for the slow axis (6240H, Cambridge Technology, Bedford, MA), achieving 15 frames per second with 500 × 500 pixels. An oil immersion objective lens (1.05 NA, UPLSAPO 30×SIR, Olympus, Waltham, MA) was used for imaging. Mineral oil (Sigma Aldrich) was used as the immersion oil due to its high transmission at 1550 nm. THG imaging required 10 nJ of 1550 nm. Three-dimensional analysis of osteocytes was performed in ImageJ 1.50i (NIH, Bethesda, MD) and Imaris 7.4.2 (Bitplane Inc., South Windsor, CT).

### Phalloidin staining

After growth in 2D or 3D, cells were washed with cold PBS after removing the culture medium. Cells were fixed with 4% PFA (Pierce^TM^, 28908) in PBS for 10 minutes and permeabilized with 0.05% Saponin (Boston BioProducts, BM-688) in PBS for 5 minutes. Cells were then incubated with phalloidin mixture (Abcam, ab176753, 1:1000) in the dark for 1 hour at room temperature with gentle agitation. Cells were washed with cold PBS three times before stained with DAPI (Invitrogen^TM^) for 15 minutes. Cells were mounted with Fluoromount-G® (SouthernBiotech) and imaged by confocal microscopy. For 10 μm tibia cryo-sections, slides were first washed in cold PBS for 5 minutes. The following steps were the same as in culture cells. Quantification of osteocyte filament density was performed in ImageJ imaging software on a blinded basis. First, cell bodies were cropped, and then multi-color images were converted to single channel (grayscale) color images. Each image was duplicated, and a binary image was created from the copied image. In the copied image, all dendrites were highlighted, and background was subtracted. Next, we used “Analyze – Set Measurements” and set “Redirect To” to the original grayscale image, followed by selecting the “Area” and “Mean grey value” functions. Finally, the “Analyze – Analyze Particles” function was used to quantify the filament density.

### Cell viability, Annexin V, and EdU assays

Ocy454 and MC3T3-E1 cells were cultured in 96-well plates at 37 °C for 7 days. From Day 1 to 7, cells were incubated each day for 1 hour at 37 °C with 10% PrestoBlue solution (PrestoBlue^TM^ Viability Reagent, Invitrogen^TM^) containing culture media (alpha-MEM supplemented with heat inactivated 10% fetal bovine serum and 1% antibiotic-antimycotic, Gibco^TM^). Absorbance was read at 570 nm (resavurin-based color change) and 600 nm (background) to calculate cell viability per the instructions of the manufacturer. To detect the apoptosis in shSp7 and shLacZ cells, we used FITC Annexin V Apoptosis Detection Kit with PI (BioLegend, 640914). We followed the manufacturer’s instructions by staining cells with FITC Annexin V and Propidium Iodide Solution for 15 minutes at room temperature in the dark. Cell apoptosis were evaluated by flow cytometry. For EdU proliferation assays, cells were labeled with 5-ethynyl-2’-deoxyuridine followed by detection by click chemistry and then flow cytometry based on the instructions of the manufacturer (ThermoFisher, C01425).

### Flow cytometry and cell sorting

For cells that were stained with phalloidin (Abcam, ab176753), cell pellets were resuspended with 2% FACS buffer (phosphate buffered saline plus 2% heat inactivated fetal bovine serum) and then filtered through round bottom tubes with cell strainer cap (Falcon®, 70 µm). For cells stained with Annexin V and EdU, cells were filtered through round bottom tubes with cell strainer cap (Falcon®, 70 µm). Flow cytometry was performed on a BD Sorp 8 Laser LSR II. For the enrichment of viable osteoblasts and osteocytes, cells were filtered through a 100 µm cell strainer to make single cell suspension. DAPI (Invitrogen^TM^) was added to cells before sorting. Dead cells, debris, doublets and triplets were excluded by FSC, SSC and DAPI. Viable tdTomato+/DAPI- cells were enriched on Sony SH800s Cell Sorter.

### Dual-luciferase reporter assay

HEK293T cells were plated in 12-well plates at the density of 50,000 cells ml^−1^. When confluent, cells were transfected using Fugene HD (Promega) with a combination of 1) pGL4 *Firefly* luciferase reporter (Promega, E6651), 2) pRL *Renilla* luciferase control reporter (Promega, E2241) and 3) FLAG-hSP7 or FLAG-hSP7^R316C^ or GFP control (VectorBuilder, EF1A as promoter, vector IDs are VB190819-1114pvu, VB190819-1115xzu and VB180924-1105fst) plasmids. Ostn_En2 sequence was synthesized with gBlocks Gene Fragments (IDT) (See sequence information in Supplementary Table 8). SacI and BglII restriction enzymes (NEB R0156S, R0144S) were used to cut both the pGL4 plasmid and the Ostn_En2 fragment. 48 hours later, cells were disassociated by adding Passive Lysis Buffer (Promega, E1910) and were gently rocking on an orbital shaker for 15 minutes. 20 µl cell lysate was transferred to 96-well black polystyrene microplates (Corning). 50 µl of Luciferase Assay Reagent II was dispensed to each well with the reagent injector. The sample plate was placed in the luminometer to measure the *Firefly* luciferase signal. 50 µl of Stop & Glo Reagent was then dispensed to each well. The plate was placed back to the luminometer to measure *Renilla* luciferase activity. Luciferase experiments were performed in biologic triplicate, and all experiments were repeated at least twice.

### RNA isolation and qRT-PCR

Total RNA was isolated from two-dimensional cultured cells using QIAshredder (QIAGEN) and PureLink RNA mini kit (Invitrogen^TM^) following the manufacturer’s instructions. Briefly, lysis buffer with 2-mercaptoethanol was added to cold PBS washed cells and collected into QIAshredder columns, then centrifuged at 15,000 g for 3 minutes. The flow-through was collected into a new tube and RNA isolation was carried out with PureLink RNA mini kit following the manufacturer’s instructions. For three-dimensional cultured cells, cells were first treated with collagenase at 37 °C for 5 minutes before isolation using the PureLink RNA mini kit. For liver RNA isolation from 6-week-old mice, RNA was extracted by tissue blender with TRIzol (Life technologies) following the manufacture’s instruction and further purification was performed with PureLink RNA mini column. cDNA was prepared with 1 µg RNA and synthesized using the Primescript RT kit (Takara Inc.). qPCR assays were performed on the StepOnePlus^TM^ Real-time PCR System (Applied Biosystems) using SYBR Green FastMix ROX (Quanta bio). *β-actin* was used as the internal control for normalization. The 2−ΔΔCt method was used to detect expression fold change for each target gene with three biological replicates.

### RNA-seq analysis

RNA sequencing was conducted using the BGISEQ500 platform (BGI, China) [74]. Briefly, RNA samples with RIN values > 8.0 were used for downstream library construction. mRNAs were isolated by PAGE, followed by adaptor ligation and RT with PCR amplification. PCR products were again purified by PAGE and dissolved in EB solution. Double stranded PCR products were heat denatured and circularized by the splint oligo sequence. The ssCir DNA was formatted as the final sequencing library and validated on bioanalyzer (Agilent 2100) prior to sequencing. The library was amplified with phi29 to general DNA nanoballs (DNBs) which were loaded into the patterned nanoarray followed by SE50 sequencing. We obtained at least 20 million uniquely-mapped reads per library sequenced. N = 2-3 biologic replicates were performed for each condition. Sequencing reads were mapped to the mouse reference genome (mm10/GRCm38) using STAR v.2.7.2b [75]. Gene expression counts were calculated using HTSeq v.0.9.1 [76] based on the Ensembl annotation file for mm10/GRCm38 (release 75). For the *in vitro* Ostn-rescue RNA-seq datasets, genes with expression counts (CPM) of lower than 0.25 in less than two samples were removed. Differential expression analysis was performed using EgdeR [77] package based on the criteria of more than two-fold change in expression value versus control and false discovery rates (FDR) <0.05. For *Sp7* knockdown and overexpression RNA-seq datasets, we first set the parameters at log_2_FC > 1 and FDR < 0.05 to identify genes with Sp7-dependent expression. We then lowered the threshold to only FDR < 0.05 (eliminating the log_2_FC cutoff) to identify genes that were counter-regulated by Sp7 for cross-referencing to Sp7 ChIP-seq datasets.

Volcano plots and heatmaps were made using EnhancedVolcano and ggplot2/tidyverse packages from Bioconductor and tidyverse (https://github.com/kevinblighe/EnhancedVolcano, https://ggplot2.tidyverse.org). Gene Ontology enrichment analysis was performed with Metascape [78] and clusterProfiler [79]. Scatter plots were made in GraphPad Prism 8.0. The degree of differential expression overlap between two transcriptomic profiles was determined by Rank-Rank Hypergeometric Overlap (RRHO and RRHO2) [80, 81]. Heatmaps generated using RRHO2 have top right (both increasing) and bottom-left (both decreasing) quadrants, representing the concordant changes, while the top left and bottom right represent discordant overlap (opposite directional overlap between datasets). For each comparison, one-sided enrichment tests were used on −log_10_(nominal *p*-values) with the default step size of 200 for each quadrant, and corrected Benjamini–Yekutieli *p*-values were calculated.

### ChIP-seq

ChIP was performed as previously described with minor modifications [27, 82]. Protein-DNA complexes were crosslinked with 1% formaldehyde; the crosslinking was quenched with 250 mM glycine. Tissues were sheared by sonication to generate a chromatin size range of 200-600 bp. Dynabeads (sheep anti-mouse IgG,11201D Life Technologies) were pre-incubated with anti-FLAG M2 antibody (F1804, Sigma-Aldrich) incubating overnight with 500 μg of chromatin. Protein-DNA crosslinks were reversed by incubating at 65 °C overnight followed by RNase and Proteinase K treatment. Samples were purified with MinElute PCR purification kit (Qiagen, 28804). The construction of ChIP-seq libraries was performed with a ThruPLEX®-FD Prep Kit (R40012, Rubicon Genomics) according to the manufacturer’s instruction. The library was sequenced on HiSeq X (Illumina) platforms.

Sequencing read quality was evaluated using FastQC (http://www.bioinformatics.babraham.ac.uk/projects/fastqc/). For both osteocyte and primary osteoblast (POB) datasets, ChIP-seq reads were aligned to the mouse genome (mm9) using Bowtie2 [83] with the parameters described previously [84]. Sp7 peaks were identified using MACS2 [85] with default parameters except for the effective genome size, which was set for mouse (mm9). The intersection between osteocyte Sp7 peaks and POB Sp7 peaks were performed with BEDTools [86]. The genomic annotation was assigned to Sp7 peaks using ChIPseeker [87]. Peaks were associated with Gene Ontology (GO) terms using the Genomic Regions Enrichment of Annotation Tools (GREAT) [88]. The assignment of target genes was performed by associating Sp7 peaks with neighboring genes using GREAT. *De novo* motif enrichment was performed within +/− 50 bps of Sp7 peak summits using DREME [29]. BigWig files from H3K27ac, H3K4me and H3K4me3 ChIP-seq experiments in IDG-SW3 cells were downloaded from the NCBI GEO database (GSE54784; [28]). The intensity of histone modifications at Ocy-specific Sp7 peaks was examined by deepTools [89].

### Serial digestion and single-cell RNA-seq

4-week old mice (n = 2 per genotype) were sacrificed and tibiae, femura, and calvariae were dissected. Soft tissue was removed through scraping, and the epiphysis was cut off. Bone marrow cells were flushed out with cold PBS using a syringe. Bones were cut into 1- to 2-mm lengths and subjected to 8 serial digestions as described previously [90, 91]. In short, bone pieces were incubated in the sequence of 15 min collagenase (0.2% of collagenase type I in isolation buffer, Worthington)/15 min EDTA solution (5 mM EDTA, 0.1% BSA in PBS)/15 min collagenase solution/15 min EDTA solution/15 min collagenase solution/30 min EDTA solution/30 min collagenase solution/30 min collagenase solution. Collagenase and EDTA solutions were prepared with RNAse inhibitor added (1:100 ratio, Lucigen, 30281). Isolation buffer is composed of 70 mM NaCl, 10 mM NaHCO_3_, 60 mM sorbitol, 30 mM KCl, 3 mM K_2_HPO_4_, 1 mM CaCl_2_, 0.1% BSA, 0.5% glucose and 25 mM HEPES. Each digestion took place in 5 ml solution in a 15 ml centrifuge tube on the thermomixer set at 35 °C/500 rpm. Bone fragments were washed in PBS between digestions. The final three collagenase fractions were collected by centrifuging the supernatant at 4 °C 300 rcf^−1^ 8 min^−1^. Cell pellets were resuspended with 2% FACS buffer containing RNase inhibitor and were filtered through a 100 µm cell strainer.

DAPI (Invitrogen^TM^) was added to cells before sorting. Dead cells, debris, doublets and triplets were excluded by FSC, SSC and DAPI. tdTomato+/DAPI- cells were sorted (Sony SH800s Cell Sorter) into a new PCR tube containing 2% FACS buffer. Single cells were encapsulated into emulsion droplets using Chromium Controller (10x Genomics). scRNA-seq libraries were constructed using Chromium Single Cell 3’ v3 Reagent Kit according to the manufacturer’s protocol. Briefly, ∼ 15,000 cells were loaded in each channel with a target output of 2,000 cells. Reverse transcription and library preparation were performed on C1000 Touch Thermal cycler with 96-Deep Well Reaction Module (Bio-Rad). Amplified cDNA and final libraries were evaluated on an Agilent BioAnalyzer using a High Sensitivity DNA Kit (Agilent Technologies). Libraries were sequenced with PE100 on the DNBSEQ^TM^ NGS technology platform (BGI, China) to reach ∼ 450 million reads.

### Single-cell sequencing read processing and analysis

Raw reads obtained from scRNA-seq experiments were demultiplexed, aligned to the mouse genome, version mm10 (with tdTomato gene inserted), and collapsed into unique molecular identifiers (UMIs) with the Cellranger toolkit (version 3.1.0, 10X Genomics). After generating digital count matrices using the count function in CellRanger, libraries were batch-corrected to achieve a similar average read count using the CellRanger aggr function. Using a cutoff of 1000 UMIs, we then performed cell type annotation iteratively through two rounds of dimensionality reduction, clustering, and removal of putative doublets across all cells. For the first level clustering, we used a modified workflow of LIGER, using integrative non-negative matrix factorization (iNMF) to limit any experiment-associated batching. Briefly, we normalized each cell by the number of UMIs, selected highly variable genes, performed iNMF across both experimental conditions (WT and *Sp7^OcyKO^*), and clustered using Louvain community detection, omitting the quantile normalization step. Visualizations were obtained and downstream trajectory algorithms were carried out on the lower-dimension embedding obtained by UMAP (Uniform Manifold Approximation and Projection).

To test for significance of proportions across all major osteoblast/osteocyte subtypes, high-quality (low percentage of mitochondrial reads) cells were used to test for differences in proportions. A two-by-two contingency table was constructed iteratively per cell type by summing all cells within and outside a defined cluster per genotype. A Barnard’s exact test was run using the Exact package in R.

Differential expression for all the major cell types were performed using a Wilcoxon rank sum test from the presto package. We removed cells with a high expression of mitochondrial genes (percentage of reads mapped to mitochondrial reads greater than 10%), removing 850 cells from the WT library and 506 from the *Sp7^OcyKO^* library. All non-osteoblast/osteocyte populations as defined by the first-round annotation were then excluded, and the modified workflow was run again to define all subpopulations. Top marker genes for all subpopulations of osteoblast/osteocytes were obtained from a Wilcoxon rank sum across all cells from the control library. The ratio of the number of osteoblast/osteocyte subtypes were determined per library by taking the number of cells within the cluster and dividing by the total number of cells obtained per library (to control for experimental differences in the number of cells captured). Differential expression between *Sp7^OcyKO^* and WT was carried out across all osteoblast/osteocyte subtypes using a Wilcoxon rank-sum test with an equal number of cells per library to ensure results were not confounded by relative changes in the number of cells with the subtype.

### Trajectory analyses – Monocle3 and velocyto

Pseudotime trajectory inference was carried out using the workflow suggested in the Monocle3 tutorial (http://cole-trapnell-lab.github.io/monocle-release/monocle3/#tutorial-1-learning-trajectories-with-monocle-3). Briefly, UMAP projections obtained for the osteoblast/osteocyte subtype analyses were used to learn a graph of cell connectivity. A root node was then defined by the most connected vertex within the Canonical Osteoblast (*Bglap3*) subtype. The pseudotime values were then displayed on the UMAP projection.

For the velocyto analyses, spliced and unspliced matrices were generated by executing the run command from the velocyto package on both libraries. Spliced and unspliced loom files were then read in using the velocyto.R package. Genes were filtered based on their expression value across all subtypes by setting the min.max.cluster.average = 0.5 for spliced and 0.05 for unspliced matrices for the control dataset and 0.8 and 0.08 for the spliced and unspliced matrices in the *Sp7^OcyKO^* dataset, respectively. The cell distance matrix was determined by creating a correlation matrix of the factors generated by the iNMF computation from LIGER. Velocities were determined using the gene-relative slopes with the following parameters (deltaT = 1, kcells = 25). Finally, velocity arrows were displayed on the UMAP embedding generated from the LIGER clustering with the following parameters (neighborhood size = 100 cells, velocity scale = ‘sqrt’ (square root),minimal cell “mass” (weighted number of cells) around each point = 0.5, number of grid points = 40).

### Heritability enrichment of cell types for human fracture risk

MAGMA (Multi-marker Analysis of GenoMic Annotation) gene annotations were carried out by running the annot command line tool across all SNPs from the hg19 human genome build. Summary statistics from fracture risk were downloaded from the Genetic Factors for Osteoporosis (GEFOS) consortium website (http://www.gefos.org/?q=content/ukbb-ebmd-gwas-data-release-2017). A z-score for all genes in the hg19 annotation was then obtained by aggregation of SNP p-values using the default settings in the MAGMA command-line tools. To test the significance of associated enrichment of fracture risk in marker genes across all major cell types, we first ran a Wilcoxon rank-sum test on all major cell types from our first round of LIGER-based clustering (aggregating all osteoblast/osteocyte subtypes together) using only cells from the wild-type mouse library. After converting mouse genes to human ones using the homolog file downloaded from the Jackson Labs website (http://www.informatics.jax.org/faq/ORTH_dload.shtml), we gathered all genes per cell type with a log-fold change greater than 0.02. The gene z-scores defined by MAGMA were then regressed against the log-fold change values per major cell type using default parameter settings from the MAGMA gene-set analysis. Bonferroni-corrected p-values and effect sizes (BETA values) were obtained for each major cell type, corresponding to the degree of enrichment of genes associated with fracture risk within the marker genes for each cell type.

### Identification of cell type-specific Sp7 target genes

10,148 Sp7-bound regions were identified as osteocyte-specific (Ocy-specific) enhancers from Sp7 ChIP-seq analysis and 6,648 genes were associated with these enhancer regions. 146 genes were counter-regulated by *Sp7* knockdown and overexpression from bulk RNA-seq analysis. 77 genes were identified as Ocy-specific Sp7 target genes when intersecting two lists of genes (6,648 and 146). Similarly, 1,733 primary osteoblast-specific (POB-specific) enhancers were identified and 1,962 genes were strongly associated with these enhancer regions. We then intersected this list of genes to the POB-specific genes derived from RNA-seq comparison between osteoblasts and chondrocytes [27], and identified 134 POB-specific Sp7 target genes.

### Sp7 target gene set enrichment analysis in DropViz dataset

To determine the degree of enrichment of cell type-specific Sp7-associated genes within the major cell types in the murine brain, a custom enrichment score was created using an empirical null distribution of gene sets from the DropViz dataset (PMID: 30096299). First, summed gene values for all transcriptionally-defined cell types were downloaded from the DropViz website (http://dropviz.org/). A Wilcoxon rank-sum test on the aggregated expression values was carried out between all major cell types in the mouse brain (Astrocytes, Choroid Plexus cells, Endothelial/Mural cells, Ependymal cells, Microglia/macrophages, Mitotic cells, Neurogenesis-associated cells, Neurons, and Oligodendrocytes/Oligo Precursor Cells (OPCs)) to define average expression differences for all genes for the major cell types. To generate a null distribution of gene set enrichment values, per cell type, the average mean difference was calculated on a randomly sampled set of 77 genes. This procedure was iterated 500 times per cell type and an empirically-defined p-value was then calculated by comparing the number of gene sets with a greater average enrichment value for that cell type than that of the Sp7-associated 77 genes, corresponding to the cell-type specific degree of enrichment for the Sp7-associated gene set. Of note, similar enrichment patterns were obtained using linear mean expression difference and fold change methods (Supplementary Fig. 8a). This same procedure was also performed on the 134 POB-specific Sp7 targets and the top 150 (ranked by p-value) marker genes of each cluster derived from mouse bone scRNA-seq dataset (Supplementary Fig. 7b, d; Supplemental Figure 7).

### Association of mature osteocyte markers with skeletal dysplasia

Mature osteocyte marker orthologs were identified in the Nosology and Classification of Genetic Skeletal Diseases [41]. Top400 mature osteocyte markers were derived from the markers identified from the cluster 6 of WT single-cell profile based on percentage of difference. Significant enrichment of mature osteocyte marker orthologs among all genes in the nosology, and within each of the skeletal dysplasia groups was calculated using RStudio (http://www.rstudio.com, V1.2.5033). The network plot was constructed using Cytoscape [92]. Each gene was colored based on percentage of difference and enriched disease groups were colored according to p-value.

### Quantification and statistical analysis

All experiments were performed at least twice. Data are expressed as means of triplicate biological repeats within a representative experiment plus/minus standard error. Statistical analyses between two groups were performed using an unpaired two-tailed Student’s t-test (Microsoft Excel). When more than group experimental groups were present, ANOVA analysis followed by post-hoc Tukey-Kramer test was performed. P-values less 0.05 were considered to be significant. Variation between groups was similar in all cases.

